# Co-cultivation is a powerful approach to produce a robust functionally designed synthetic consortium as a live biotherapeutic product (LBP)

**DOI:** 10.1101/2021.10.13.464188

**Authors:** Fabienne Kurt, Gabriel E. Leventhal, Marianne R. Spalinger, Laura Anthamatten, Philipp Rogalla von Bieberstein, Gerhard Rogler, Christophe Lacroix, Tomas de Wouters

**Author notes:** Senior authors (Christophe Lacroix, Tomas de Wouters).

## Abstract

The successes of fecal microbiota transplants (FMT) have provided the necessary proof-of-concept for microbiome therapeutics. Because of the many risks and uncertainties associated with feces-based therapies, defined microbial consortia that modify the microbiome in a targeted manner have emerged as a promising safer alternative to FMT. The development of such live biotherapeutic products has important challenges, including the selection of appropriate strains and the production of the consortia at scale. Here, we report on an ecology and biotechnology-based approach to microbial consortium design that overcomes these issues. We designed a nine-strain consortium that emulates the central metabolic pathways of carbohydrate fermentation in the healthy human gut microbiota. We show that continuous co-culturing the bacteria produce a stable consortium whose activity is distinct from an equivalent mix of individually cultured strains. Further, we showed that our function-based consortium is as effective as FMT in counteracting dysbiosis in a dextran sodium sulfate mouse model of acute colitis. We propose that combining a bottom-up functional design with continuous co-cultivation is a powerful strategy to produce robust, functionally designed synthetic consortia for therapeutic use.

---

Intestinal microbiome based live biotherapeutic products (LBPs) are emerging as a novel modality to treat a large number of chronic diseases^1,2^. Tire therapeutic objective is to induce a targeted modulation of the intestinal microbiota by administering live microorganisms to reverse dysbiosis and promote recovery^3,4^. However, how to design and produce such LBPs and robustly achieve intestinal microbiota modulation is a yet unsolved technological and biological challenge. Fecal microbiota transfer (FMT) from healthy donors is currently the most successful method of restoring diseased microbiota to a healthy state^5^. However, it is not fully understood what the specific properties of a healthy microbiota are and how these might promote recovery^6,7^, resulting in variable efficacy of FMT for different diseases^7-11^. Moreover, the potential for adverse events with FMT raises critical safety concerns^12^.

Defined bacterial consortia that are well-characterized represent a safer and more controlled alternative to FMT^13,14^ Because a healthy microbiota typically contains hundreds of strains, simply recreating the full taxonomic diversity in a defined product is intangible. The complexity can, however, be reduced by removing functionally redundant strains^15-17^ and focusing on those microbial functions that are key for the specific therapeutic target. While intuitive in principle, translating this functional concept into a defined consortium is not straightforward because most of the intestinal microbiota is still uncultured and uncharacterized^18^. Here, we present a technological solution to designing and coculturing multi-strain consortia based on the principle of ‘division of labor’. Microbial consortia in which the overall function is divided among bacteria are often more robust and productive than would be expected from the physiology of the individual strains^19-23^. Examples include simplified trophic consortia used in wastewater treatment^24^, biofuel production^25-27^, and food production^28^. Nevertheless, this principle has not yet been applied to designing intestinal LBPs. We hypothesized that division of labor increases consortium stability and robustness, thus enabling the resulting LBP to match the therapeutic effect of FMT. We used our approach to design a consortium of nine anaerobic intestinal bacteria that cover the essential elements of carbohydrate metabolism in the large intestine^29,30^. Using continuous co-cultured fermentations, we confirmed the establishment of a compositional and metabolic equilibrium with complete carbohydrate fermentation akin to a healthy intestinal microbiome. Importantly, co-culturing affected the phenotypic state of the strains in such a way that the co-cultured consortium (CC) was different than an equivalent mix of identical but individually cultivated bacterial strains. Finally, the CC matched *in vivo* efficacy of FMT in a mouse model of acute Dextran Sulfate Sodium (DSS) colitis, while the simple mix of strains did not. Thus, our approach offers a robust blueprint for the design and production of consortium LBPs.

## Results

### The joint metabolic activity of nine strains fully converts complex carbohydrate substrates into end metabolites without accumulating intermediate products

We aimed to develop a simplified bacterial consortium that recapitulates the most central function of the microbiota in a healthy intestine—carbohydrate fermentation. On one hand, bacteria gain energy from breaking down complex carbohydrates for growth and maintenance. On the other hand, fermentation products interact with host physiology. The key end products of carbohydrate metabolism are short-chain fatty acids (SOFA) that can be beneficial to the host^31,32^. However, the accumulation of intermediate breakdown products due to incomplete fermentation can be detrimental^32-34^. This multi-step conversion of complex carbohydrates through intermediate products into end products in the gut is typically distributed across different bacterial species in a multi-level trophic cascade. While there are several trophic paths to convert one metabolite into another^35,36^, we here propose a defined set of 13 metabolic reactions deemed as ‘essential’ based on the reviewed literature. These reactions reflect the baseline requirement for the stabilization of the gut microbiome as an ecosystem (Figure 1a). Primary **A’** reactions cover the conversion of complex fibers, starches, and sugars into either intermediate (formate, lactate, succinate, acetate) or end products (acetate, butyrate, propionate). Primary degraders perform these reactions using specific mechanisms such as extracellular enzymes to de grade and import these primary substrates. Secondary ‘B’ reactions cover the conversion of intermediate metabolites into the end metabolites. These intermediate products can inhibit the growth of some bacte ia^37^ and have been associated with different diseases^32,33,38^. Finally, ‘C’ reactions consume gases like hydrogen and oxygen that typically have inhibitory effects at high concentrations by disrupting redox balance or imposing oxidative stress^39^. With this characterization of the trophic cascade at hand, we next proceeded to design a consortium to implement this functional sequence of reactions.

**Figure 1:**
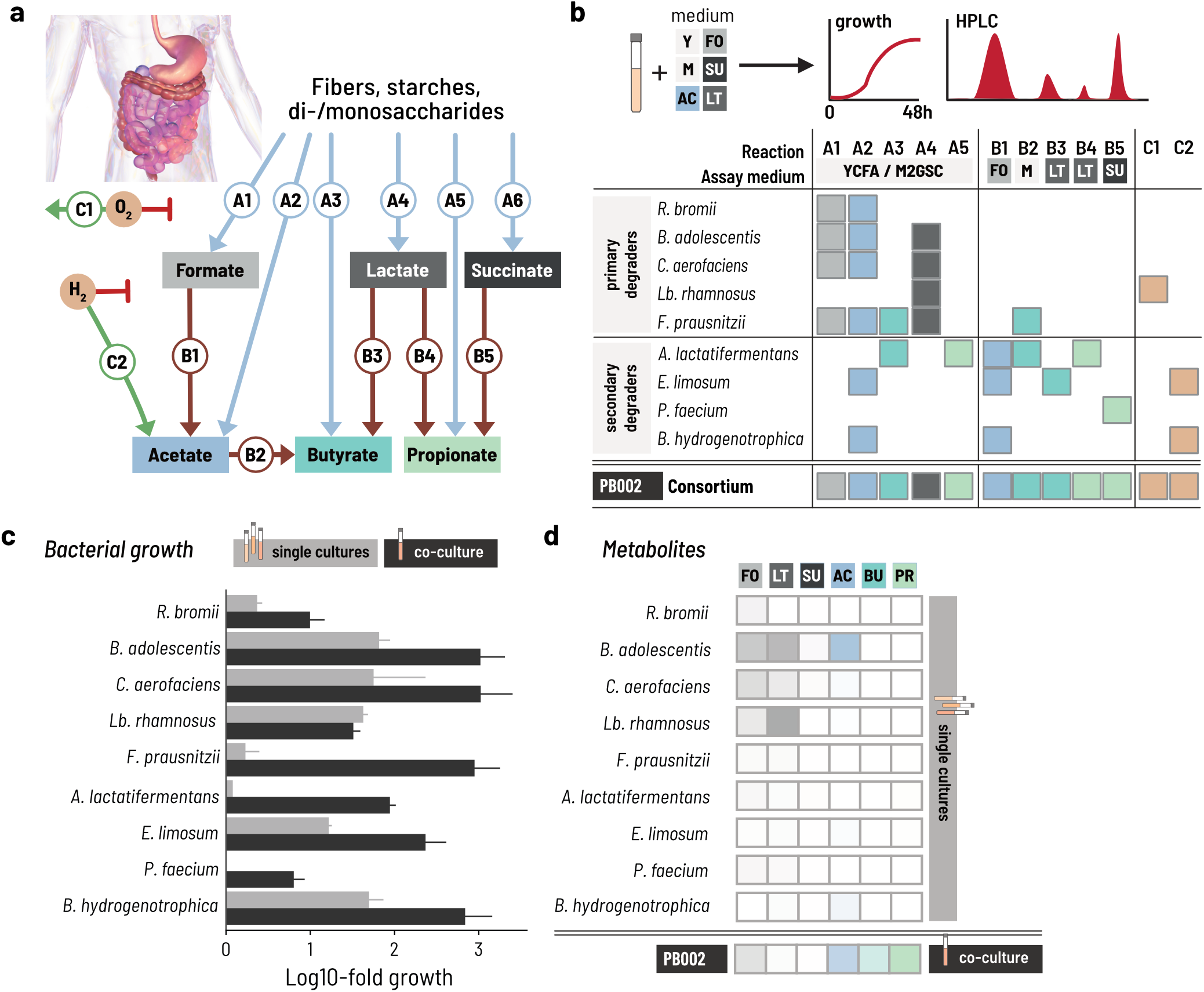
A minimal intestinal microbiome of nine strains performs complete carbohydrate fermentation, **a**. The trophic cascade of essential metabolic reactions ferments carbohydrates into the end metabolites acetate, propionate, and butyrate. Formate, lactate, and succinate are common intermediate metabolites that might be produced and secreted. ‘A’ reactions convert primary substrates to intermediate or end metabolites, and ‘B’ reactions convert intermediate metabolites. ‘C’ reactions consume oxygen and hydrogen that can inhibit fermentative activity, **b**. The combined metabolic activity of nine selected species covers all essential reactions for transforming primary substrates and intermediate metabolites into end metabolites. We assayed metabolite consumption and production by growing strains in media containing either sugar and/or starches or specific intermediate metabolites as carbon sources. Five primary degraders produce intermediate metabolites and acetate from primary substrates. Four secondary degraders consume intermediates and produce end metabolites. Hydrogen and oxygen metabolism were attributed to strains using published data. **c**. Most of the strains had equivalent or larger growth in co-culture than in single culture in the PBMF009 medium. Log fold change of single strains (in grey) and co-cultured strains (in black) as determined by qPCR for three replicate experiments with three technical replicates from time point 0 h and 48 h with S0. The data are normalized by the 16S rRNA gene copy number of each strain. *P. faecium* and A. *lactatifermentans were* not able to grow in single culture in the PBMF009 medium, **d**. Primary degraders predominantly produced formate and lactate in single culture, except for *B. adolescentis* that also produced acetate. All four secondary degraders did not produce more than 10 mM of total metabolites in single culture. The co-culture of all nine strains, however, produced more than 74.3 ± 4.5 mM of metabolites, thereof predominantly end metabolites, acetate, propionate, and butyrate. Y: YCFA, M: M2GSC, F0: formate, LT: lactate, SU: succinate, AC: acetate, BU: butyrate, PR: propionate. Intestine image adapted from ref [40].

We selected a set of bacterial strains to fully cover these essential reactions and hypothesized that the resulting consortium would establish a trophic reaction cascade that mimics the carbohydrate metabolism in a healthy gut microbiome. To this end, we profiled a panel of human intestinal isolates for A and B reactions by testing their metabolic activity in anaerobic batch fermentations. For A reactions, we assessed growth after 48 h on media containing sugars and starches as carbon sources (M2GSC, YCFA). For B reactions, we assessed growth on media containing the respective intermediate organic acid as a carbon source after 7 days to account for the slow utilization rates we observed under these conditions. We assigned C reactions to the strains using published data: oxygen reduction was assigned to strains reported to grow under aerobic or microaerophilic conditions and hydrogen consumption to acetogenic or methanogenic gut bacteria^41-44^. Using this approach, we selected a minimal set of nine strains that cover all essential reactions of the carbohydrate metabolism except for succinate production (A6) and subsequently refer to this consortium as PB002 (Figure 1b, Supplementary Figure S1). We excluded succinate producing *Bacteroides* species due to its association with inflammatory outcomes in our models. Interestingly, genome-based *in silico* predictions of the metabolic capabilities of the nine strains uncovered only a small fraction of their actual measured capabilities (Supplementary Table S4), confirming that *in vitro* phenotypic characterization of strains is required.

Tie individual strain assays show that each has different requirements for growth. However, because we chose the strains based on their trophic interactions, we hypothesized that the metabolic interactions within the consortium would allow all nine strains to grow even in a medium that did not explicitly accommodate all their specific requirements. Rather, we designed the medium to contain a minimal amount of undefined ingredients that are typically used to cover unknown nutrient requirements. The medium—PBMF009—contained four different primary substrates—disaccharides, fructooligosaccharides, resistant starch, and soluble starch— and was free of rumen fluid and meat extract but did contain yeast extract (Supplementary Table S1). We then measured the growth and metabolic activity of the nine strains individually and in co-culture in PBMF009 in naerobic batch incubations for 48 h. We quantified the growth of each strain by qPCR with genus-specific primers normalizing for gene copy number (Supplementary Table S2) and measured the metabolite profiles using high-performance liquid chromatography with refractive index (HPLC-RI).

All primary degraders showed equal or increased growth when co-cultured compared to monocultures (Figure 1c, Supplementary Figure S2). In monocultures, three out of the five primary degraders, *Collinsella aerofiaciens, Bifidobacterium adolescentis*, and *Lactobacillus rhamnosus* grew more than 1.5 orders of magnitude after a 1% v/v inoculation, confirming that these primary degraders can utilize elements of PBMF009 as a growth substrate. Notably, these primary degraders grew equally well or better in co-culture than in monoculture. The remaining two primary degraders, *Ruminococcus bromii* and *Faecalibacterium prausnitzii*, grew poorly over the 48 h in monoculture, but their growth was boosted in co-culture, with *E prausnitzii* achieving nearly a three-log growth. Because primary degraders are expected to compete for carbon sources when co-cultured, the superior growth in co-culture suggests that the strains are utilizing different components of the medium as their primary energy source. Also, the presence of the other strains improves the ability to extract energy for growth from the medium beyond what is possible in monoculture.

The growth of secondary degraders was equally boosted in co-culture compared to monoculture (Figure 1c, Supplementary Figure S2). In monoculture, *Eubacterium limosum* and *Blautia hydrogenotrophica* showed intermediate regrowth between 10-to-30-fold that increased to over 100-fold in co-culture. *Phascolarc-tobacterium faecium* and *Anaerotignum lactatifiermentans* marginally grew in monoculture. Their growth was enhanced in co-culture, suggesting that the compounds required for their growth were produced by the other strains in the consortium.

The metabolite profile of the co-culture suggested a successful establishment of the designed trophic network (Figure 1d, Supplementary Figure S3). In monoculture, all primary degraders produced mainly formate or lactate between 5-15 mM, and *B. adolescents* additionally produced acetate above 30 mM. The secondary degraders did not produce meaningful amounts of metabolites in line with their poor growth in monoculture. In co-culture, we did not detect notable levels of lactate at 24 or 48 h. While formate reached moderate levels up to 5 mM at 24 h it decreased again by 48 h. This suggests that the lactate and formate that was produced by the primary degraders was subsequently metabolized by the secondary consumers, resulting in the production of propionate, butyrate, and acetate.

### An initial fed-batch fermentation using ‘controlled strain ratios’ promotes the balanced growth and complete carbohydrate metabolism

Given that co-culturing improved the growth of all strains when compared to equivalent monocultures, we next tested whether strain coexistence and trophic interactions were robustly maintained in a scalable bioreactor setup. To this end, we mixed all strains together at equal volumes in condition-controlled stirred-tank bioreactors run in batch mode at physiologically relevant conditions (pH = 6.0, 37 °C, anaerobiosis) for 48 h (Figure 2a). Starting 12 h post inoculation, we monitored optical density (OD) and metabolite concentrations every 3 h, and strain composition for a subset of time points.

**Figure 2:**
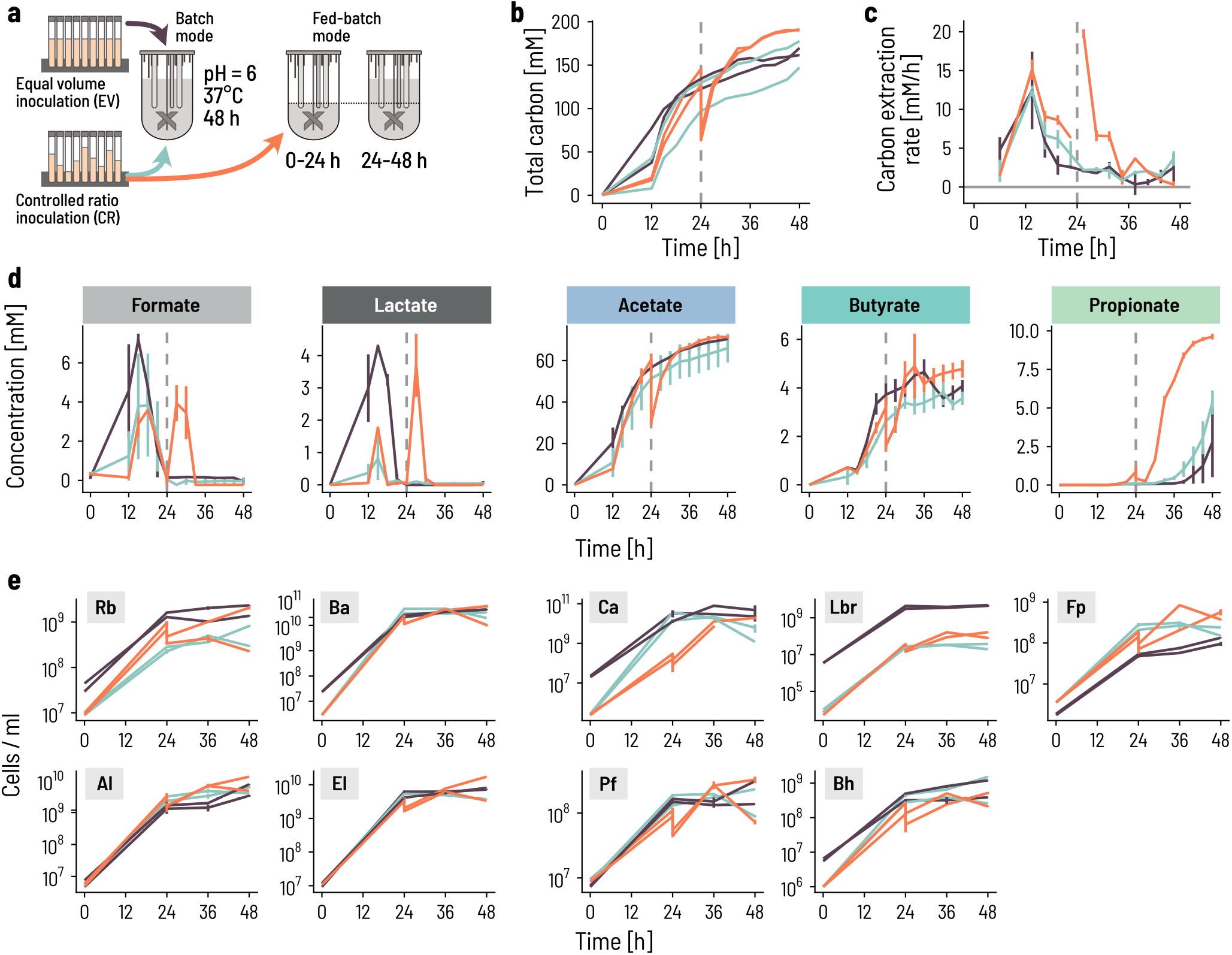
Fed-batch and controlled inoculation ratios of strains promote the successional colonization of bioreactors, **a**. We compared two adjustments to the fermentation strategy: using either a batch or a fed-batch setup over 48 h with two different inoculation ratios, ‘equal volumes’(EV; black lines) or‘controlled ratio’ (CR; green or orange lines). Batch fermentations were inoculated at 0.33% v/vand fed-batch bioreactors at 0.67% v/vto balance the doubling of medium during fed-batch fermentation. The volume was doubled by adding fresh medium after 24 h in fed-batch mode. **b**. The total amount of metabolites produced, expressed in carbon mol concentration, increases steadily in batch mode with either inoculation ratios (EV: black, CR: green). There is a visible 2-fold drop at 24 h in fed-batch mode as a result of medium addition (orange). Each of the three conditions were run in duplicate. Metabolites were quantified by HPLC-RI analysis. Total carbon production was calculated by summing the C-mol of all measured metabolites. The C-mol concentration was calculated by the product of the metabolite molar concentration and the respective number of carbon atoms, **c**. We computed the rate of carbon extraction as the empirical difference of the C-mol concentration during the last measurement period and expressed per unit time (hour). The lines show the mean of, and the vertical segments show the spread between the duplicates, **d**. Intermediate (formate, lactate) and end (acetate, butyrate, propionate) metabolites are dynamic during the 48 h fermentation. Each line shows the mean of, and the vertical segments the spread between the duplicates, **e**. The growth of the nine strains differed between the strategies. Fermenting in fed-batch mode increased the subseguent growth of the secondary degraders in the second stage. Cell counts were determined by qPCR, whereby each time point was measured in triplicates. The lines show the median and the vertical segments show the min and max of the triplicates. The measured gene counts are normalized by the 16S rRNA gene copy number of each strain.

Despite a rapid initial increase in metabolic activity, carbon conversion of the co-culture was not complete by 48 h. We computed the total amount of measured metabolites expressed in mols of carbon (C-mols) as a marker of overall consortium activity. C-mols increased sharply between 12-16 h (Figure 2b,c), mainly as a result of the production of acetate, lactate, and formate (Figure 2d), in line with dominant metabolic activity of fast-growing primary degraders. After this initial spike, formate and lactate concentrations dropped to zero, suggesting a delayed onset of growth and activity of secondary degraders and the successful establishment of the full metabolic cascade. Subsequently, C-mols increased slowly but did not level off before 48 h, implying that the co-culture was not able to fully convert the available energy in the fermentation medium.

To accelerate the establishment of an equilibrated coculture, we adjusted the inoculation ratios to limit the growth of fast-growers and promote the growth of slow-growing secondary degraders (Figure 2a). The resulting ‘controlled strain ratios’ were based on the natural abundance in the specific host microbiota from which we isolated seven of the nine strains. Additionally, we diluted *Lb. rhamnosus* ten-fold and increased *E. limosum, P.faecium*, and *A. lactatifermentans* roughly five-fold (Supplementary Table S5).

Inoculating at ‘controlled strain ratio’ and fermenting in batch mode did not improve the co-culture efficiency, and resulted in similar overall activity profiles, as indicated by similar amounts of produced C-mol up to 48 h post inoculation (Figure 2b,c). As expected, we measured less formate and lactate compared to equal volumes at 12 h, which also translated into less butyrate beyond 24 h. The adjusted inoculation ratios were successful in reducing the cell numbers of the dominant *R. bromii* and *Lb. rhamnosus* 10 and 100-fold, respectively However, this did not impact *B. adolescentis* and *C. aerofaciens* over the first 12 h (Figure 2e). The growth of these latter two strains was impaired beyond 24 h compared to the equal volumes. Apart from a small boost in growth for A. *lactatifermentans*, we did not observe any benefit for the secondary degraders. This suggested to us that while setting a cap on the starting inoculation levels of fast growers such as *Lb. rhamnosus* can reduce their early dominance, adjusting the inoculation ratios alone is not sufficient to boost growth and activity of the secondary degraders. We thus implemented a two-step fed-batch mode to limit the growth of primary degraders and further support secondary degraders. We started the co-culture with half the volume of fermentation medium and supplied the second half after 24 h.

The fed-batch strategy was successful at enabling the growth of secondary degraders and therefore promoting trophic succession. Tie overall behavior of the co-culture in terms of metabolite profile and strain composition was practically identical to the controlled ratios until 24 h, implying that the reduction in fermentation volume over the first 24 h had little effect. Adding the remaining culture medium at 24 h brought an additional boost to activity and growth. The carbon conversion rate spiked immediately after feeding and subsequently tapered off to zero by 48 h (Figure 2c). Adding the fed-batch step thus enabled the co-culture to rapidly and completely metabolize the available carbon in the medium and reach an overall plateau (Figure 2b). A ‘second phase’ growth was visible in all primary degraders, including a large boost for *F. prausnitzii*. This strain reached almost two-fold higher cell numbers in fed-batch mode than in regular batch mode. The growth of the secondary degraders was also boosted immediately after feeding. This was especially apparent for *A. lactatifermentans* and *P faecium*, which showed strong growth after 24 h and an overall stronger production of propionate. Thus, the two-stage fed-batch mode allowed the slower growing secondary degraders to catch up with the fast growers during the first stage and enable the co-culture to enter the second stage in a more synchronized manner.

### Continuous co-cultivation of the consortium results in stable strain ratios, with high cell yields and end metabolite production

The differences in outcome between the inoculation strategies imply an important influence of initial conditions on co-culture dynamics and activity. Nevertheless, we hypothesized that if all strains were retained during the initial phase, then the trophic interactions would dictate the final composition of the culture and lead to a specific equilibrium. To determine whether such a stable compositional equilibrium exists and what its strain composition is, we switched the fermentation to continuous mode after the initial fed-batch mode. We chose a high mean retention time of 50 h (D = 0.02 h ^1^) to mimic the slow transit in the human colon and to limit washout of slow growers. We measured the metabolite profile every 24 h for two weeks and quantified the community composition on a subset of days (Figure 3a). To test whether different inoculum ratios lead to the same or alternative compositional equilibria throughout the continuous fermentation period, we inoculated the batches using both the controlled strain ratios (CR) and the equal volumes (EV).

**Figure 3:**
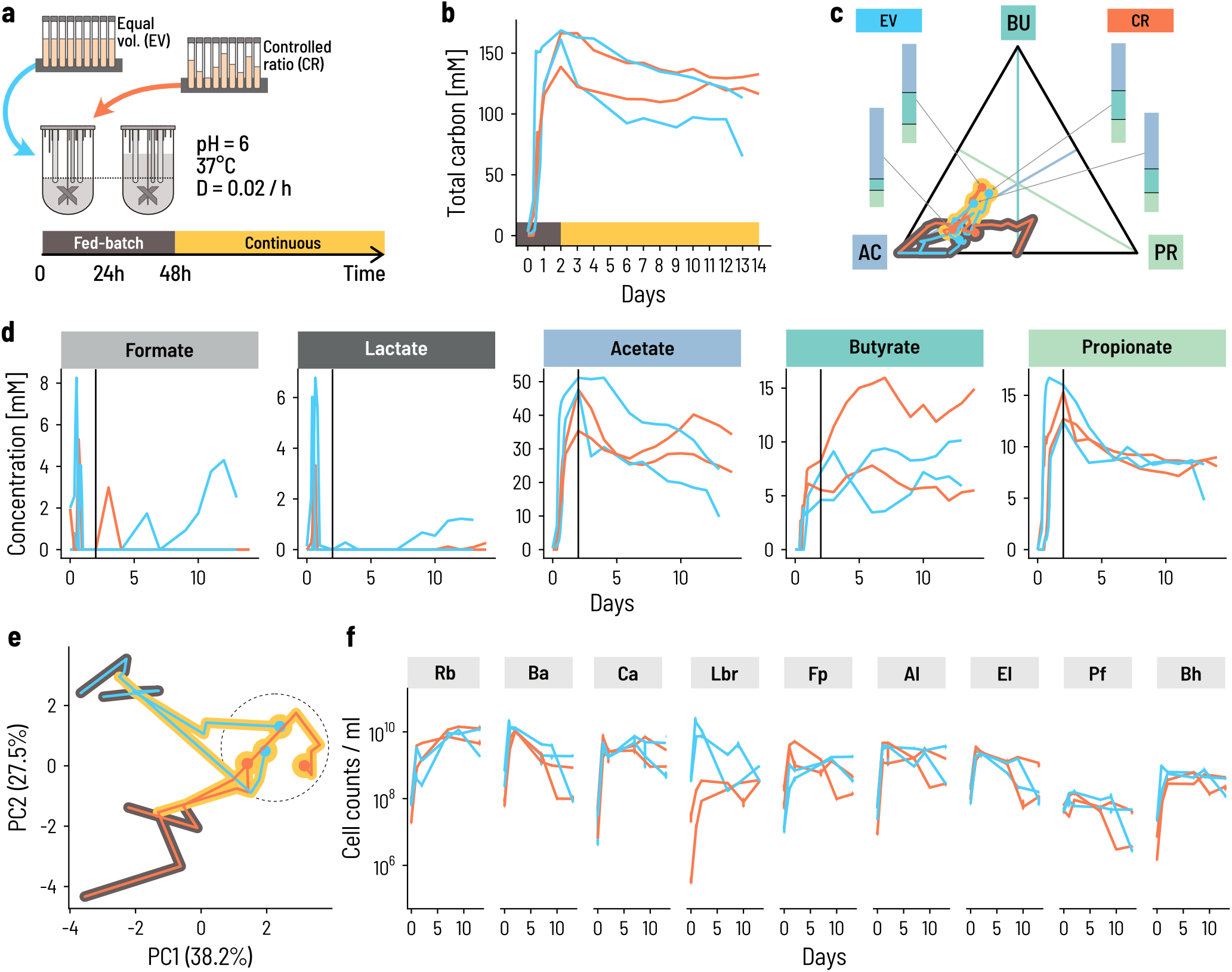
Continuous co-cultivation directs the consortium into a characteristic metabolic and compositional equilibrium, **a**. Four bioreactors were inoculated at 0.67% v/v at either equal volumes (blue lines) or controlled ratios (orange lines). Fed-batch fermentation was carried out for the first 48 h, whereby the fermentation volume was doubled after 24 h of fermentation by adding fresh PBMF009 medium. After 48 h, the system was set to continuous mode. Fermentation conditions were chosen to reflect conditions of the human colon, at 37 °C, pH 6.0 and a mean retention time of 50 h, or 0 = 0.02 / h. **b**. The total amount of carbon in the measured metabolites increases rapidly during the fed-batch phase and subsequently levels off during the continuous phase, **c**. The relative molar concentration of the end metabolites changes throughout the experiment but reaches a common endpoint. The position on the simplex shows the relative proportion of acetate (AC), butyrate (BU), and propionate (PR). The closer a point is to a corner, the more of the respective metabolite is in the sample. Circles indicate the endpoints of the experiment and for each one the stacked bar shows the relative proportions. The black and yellow background of the lines indicates the fed-batch and continuous phase, respectively, **d**. The concentration of metabolites changes over the course of the experiment. Succinate was below detection in all samples and is not shown, **e**. The strain composition at sampling points throughout the experiment evolves from the inoculum composition to a common equilibrium composition (dashed circle). Principal components (PC) are computed from the center log-ratio transformed relative cell concentrations. The circles show the endpoint of the fermentations, and the black and yellow background corresponds to the fed-batch and continuous phase, respectively, **f**. Cell concentrations for each of the nine strains are dynamic through time. Rb: R. *bromii;* Ba: *B. adolescentis;* Ca: C. *aerofaciens;* Lbr: Lb. *rhamnosus;* Fp: F. *prausnitzii;* Al: *A. lactatifermentans;* El: E. *limosum;* Pf: P. *faecium;* Bh: *B. hydrogenotrophica*. Each sampling point is the median of three technical qPCR replicates and the error bars show the minimum and maximum measurements.

The overall metabolic activity was maintained at high levels after switching from fed-batch to continuous mode. The total amount of carbon in the measured metabolites decreased slightly after changing the operation mode and approached an apparent stable level after approximately 10 days for the CR inoculum (Figure 3b). The relative contribution of acetate, butyrate, and propionate to C-mols varied between replicate reactors during the fed-batch phase, but eventually converged to a common composition with 56% acetate, 24% butyrate, and 20% propionate on average (Figure 3c). Propionate levels were extremely consistent and approached a mean of 8 mM across all four reactors (Figure 3d). Acetate and butyrate levels were more variable. In all reactors, acetate was highest at the end of the fed-batch phase and decreased during the continuous phase, while butyrate levels were maintained or even increased. This is consistent with the notion that acetate is both a general marker for metabolic activity and an intermediate compound in butyrate production. Acetate stabilized after *5* days in the CR reactors but continued to decrease through the full experiment for the EV reactors, implying that the CR inoculum enabled the consortium to approach its metabolic equilibrium more rapidly. In this equilibrium, butyrate and acetate were strongly anticorrelated (Pearson’s r = −0.83, p = 0.00485 and r = −0.81, p = 0.00113, respectively), suggesting that this fluctuation arises from the conversion of acetate into butyrate.

In all four CR and EV bioreactors, all nine strains persisted during continuous planktonic fermentation, although their relative abundance changed after switching to continuous mode. The initial reactor compositions were different because of the different inoculation strategies but converged to a common state on day 13 (Figure 3e). At the end of the fed-batch phase, the dominant primary degraders shifted from *B. adolescentis* and *C. aerofaciens—*and *Lb. rhamnosus* in the case of EV—to *R. bromii* that comprised 65% and 50% of the consortium at the end of the experiment for CR and EV, respectively (Figure 3f). *Lb. rhamnosus* initially differed between CR and EV, but converged to a common equilibrium level in CR and EV. All other strains had dynamics that did not strongly differ between CR and EV.

These data suggest that the combination of trophic design principles for co-cultures and the optimization of process conditions gives rise to a stable, reproducible consortium both in terms of metabolic output and composition. To validate this reproducibility, we performed four additional repetitions of the fed-batch and continuous fermentations with CR for extended periods of up to 92 days (Supplementary Figure S4). Despite early transient differences, the composition of the community consistently reached a state of equilibrium after ca. 10 days of continuous cultivation. We therefore concluded that this state of equilibrium is intrinsic to the trophic structure of the consortium in the context of the biotechnological cultivation process.

### The consortium state at the end of co-cultivation is not recapitulated by a mix of strains at the same ratios

We next assessed whether the PB002 consortium at equilibrium has phenotypic characteristics beyond the equilibrium ratios of the nine strains. Because of the trophic interactions that underlie the consortium’s bottom-up design, we hypothesized that the stable co-existence of all nine strains implies a physiological synchronicity between them. If this were the case, merely mixing the strains together at equilibrium ratios would not mimic this synchronization of the strains. To test this hypothesis, we performed batch fermentations in serum bottles and compared the regrowth of the CC to individually grown single strains mixed at the same strain ratio. To produce the two different inocula, the strains were cultured either individually in Hungate tubes in their preferred growth medium or co-cultured in a bioreactor until the previously defined equilibrium state was reached. The individually cultured strains were mixed to match the absolute abundance of the CC at equilibrium to form an ‘artificial mixture’ (AM). Batch fermentations were performed in triplicates for 48 h using 3x buffered PBMF009 medium. We measured the consortium composition by qPCR and the metabolites by HPLC-RI every 12 h (Figure 4a).

**Figure 4:**
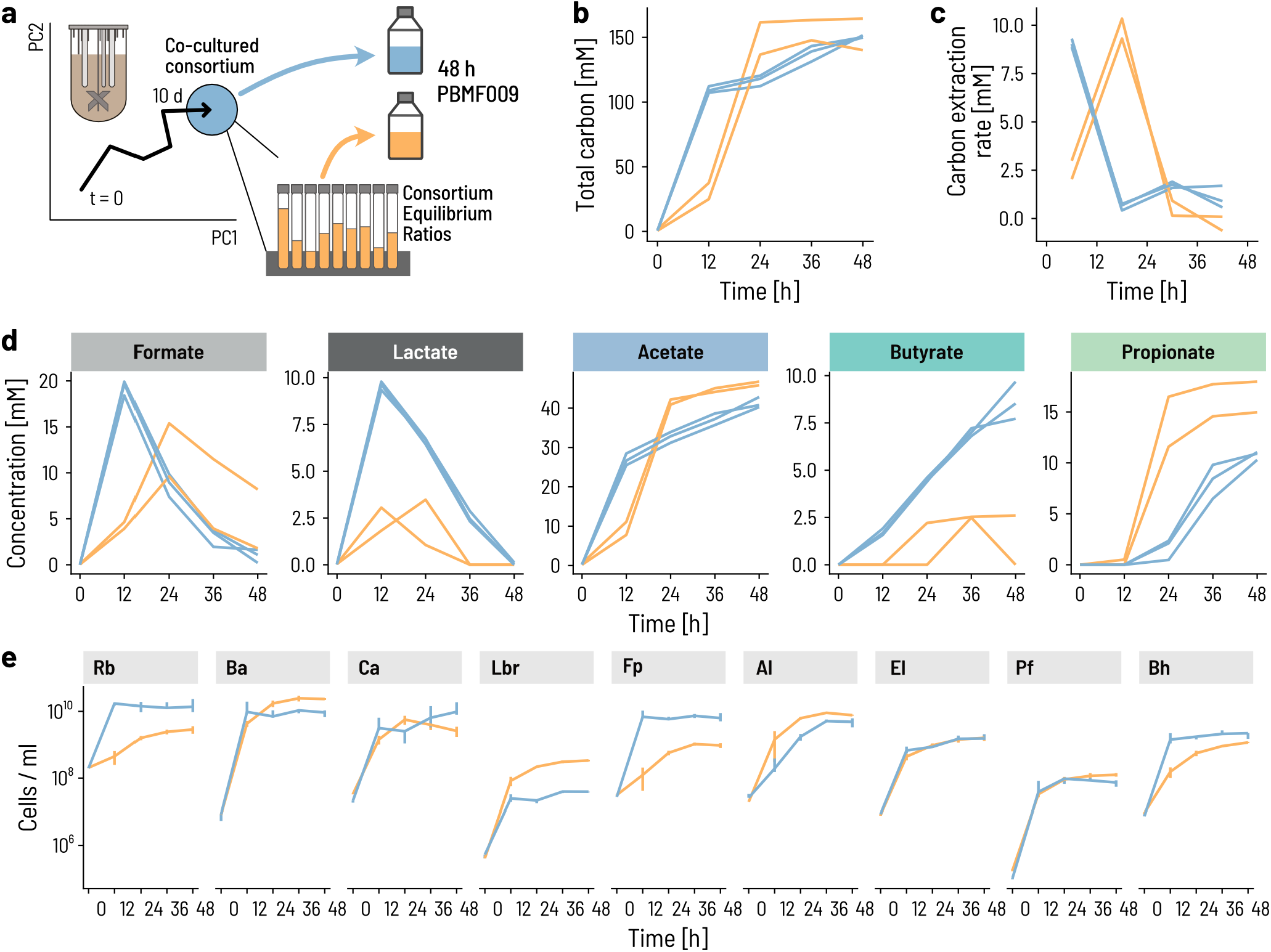
The onset of growth and activity occurs sooner for the consortium than a mix of strains, **a**. We compared the growth and metabolic activity of the co-cultured consortium to that of a mix of the same nine strains pooled together to imitate the cell counts of the individual strains at equilibrium. **b**,**c**. The total amount of metabolites produced, expressed in C-mol concentration increases more rapidly in the consortium than in the mix of strains, **d**. Formate and lactate accumulate earlier and to higher concentrations in the consortium. Acetate levels follow the same dynamics as total carbon. The consortium produces more butyrate, while the mix of strains produces more propionate, **e**. Cell counts of each of the nine strains were measured using qPCR. Strains with a slower growth rate in the strain mix grew earlier in the consortium.

The CC cultures were already metabolically active with an early peak of overall carbon extraction rate between 0 and 12 h, whereas peak extraction for AM occurred between 12 and 24 h (Figure 4b,c). The activity of AM surpassed the consortium at 24 h, mostly because of increased acetate production likely by the fast-growing primary degraders (Figure 4d). Formate and lactate concentrations were higher in the CC than the AM after 12 h, with higher butyrate production in the CC samples. In contrast, higher propionate levels in the AM suggested that the absence of a lactate buildup in these samples was likely due to a rapid conversion of lactate into propionate. These metabolic patterns were consistent with strong differences in strain growth between the CC and the AM. Most strikingly, those strains for which we observed a ‘conditioning’ during the continuous fermentation such as *R. bromii* and *F. prausnitzii* grew without delay in the CC samples but not in the AM samples (Figure 4e). In addition, the previously observed fast growers *Lb. rhamnosus, B. adolescentis*, and *C. aerofaciens* were dampened in the CC versus the AM. These results confirm our hypothesis that the physiological state of the individual strains at co-culture equilibrium is adapted to the ‘consortium state’ and cannot be replicated by merely mixing the strains at the equivalent ratios.

### The PB002 consortium promotes recovery from acute DSS colitis in mice as efficiently as FMT

We postulated that the same kind of ‘physiological conditioning’ of the strains through co-cultivation would occur during ‘natural co-cultivation’ in the human intestine. We thus hypothesized that a co-culture of PB002 would better match the therapeutic effects of FMT than a mix of individually cultivated strains. We tested this through two independent experiments using the acute DSS-induced colitis mouse model (Figure 5a). Specifically, we compared treatment with our CC to FMT from a healthy human donor. To tease apart the effect of co-culturing from the individual strains or the produced metabolites only, we also treated mice with a mix of individually cultured consortium strains (SM), the conditioned medium harvested from the continuous fermentation (CM), and phosphate-buffered saline as a control (PBS).

**Figure 5:**
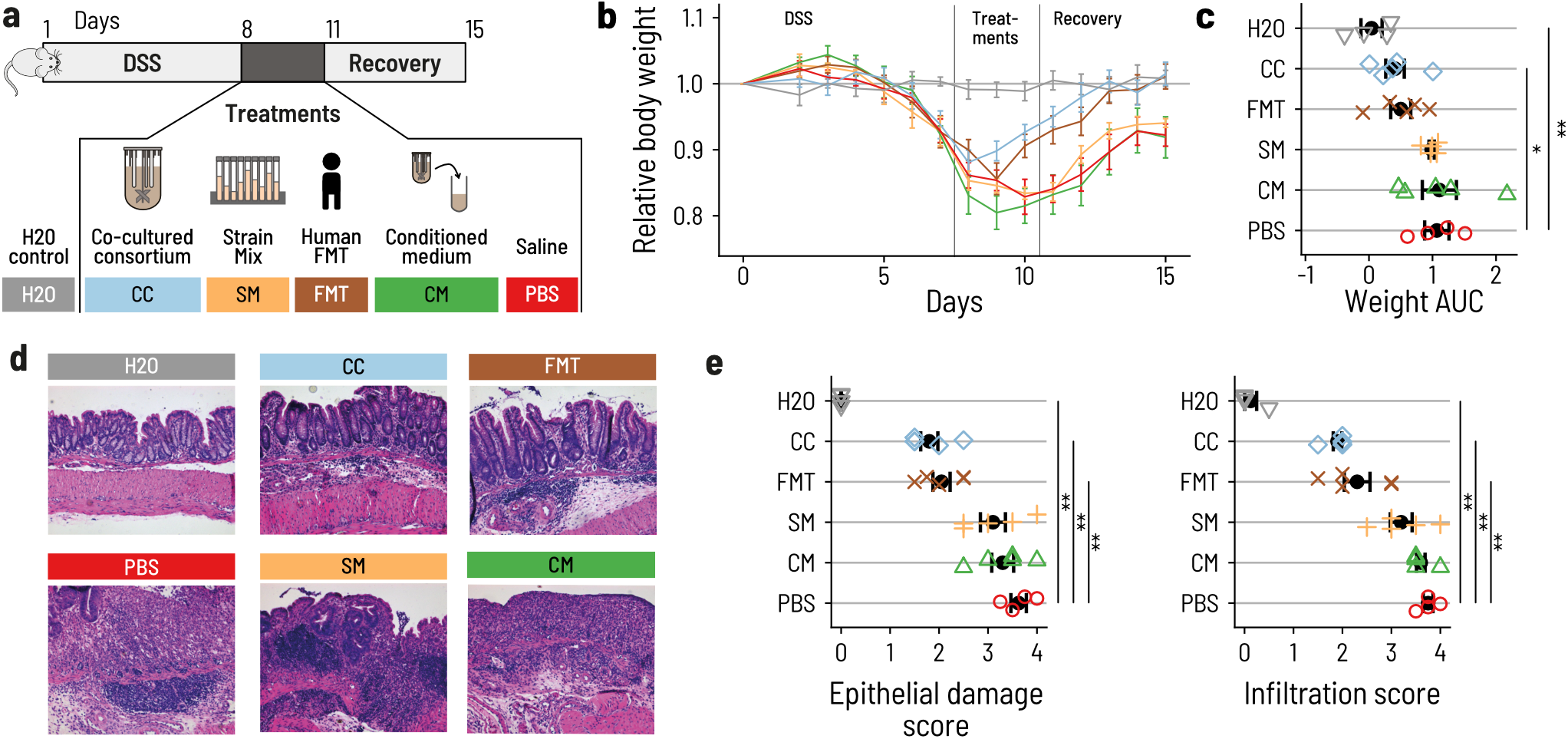
PB002 accelerates recovery after DSS-induced colitis in mice. **a**. Acute colitis was induced in female C57BL/6 mice by supplementing the drinking water with OSS for seven days (from day 1 to day 8). At day 8, mice were switched back to normal drinking water. Mice were treated once a day by 200 pL oral gavage on days 8, 9, and 10 with the co-cultured consortium (CC, blue), the non-co-cultivated strain mix(SM, orange), the conditioned medium from the continuous fermentation (CM, green), or with FMT from a healthy human donor (FMT, brown). The control group was given normal water throughout the whole experiment (H20, grey), and the OSS control group was gavaged with phosphate-buffered saline (OSS, red). Mice were euthanized at day 16. N = A to 5 per group, **b**. Mice treated with CC or FMT regained body weight more rapidly than mice that received the other treatments. Error bars are the standard errors, **c**. Area under the curve (AUC) of the daily relative body weight for each mouse. Only CC-treated mice had an AUC that was significantly lower than the OSS control mice (linear model, *β* = 0.65, *p* = 0.030). The error bars show the confidence interval for the mean (1.96 x SEM). d. Representative light micrographs of large intestine sections at the time of euthanasia (H&E staining, 10 X magnification). CC and FMT showed structural recovery of the epithelium comparable to the control group. Mice that received CM, SM, or PBS showed a substantial degradation and inflammation of the cecal epithelium, **e**. Histological assessment of the epithelial damage and infiltration in the distal colon. Treatment with CC and FMT showed reduced epithelial damage and infiltration compared to PBS (linear model, *p <* 10^−4^ for all marked comparisons)

A drastic body weight loss was observed in all DSS-treated mice. Animals treated with the CC or FMT regained weight by day 10 and fully reached their initial body weight by day 14. Body weight recovery was much faster in mice that received CC or FMT compared to mice that received PBS, SM, or CM, in which body weight only increased after day 12 (Figure 5b, Supplementary Figure S5). Overall, mice that received the CC lost significantly less body weight throughout the experiment than PBS treated mice (Figure 5c). To test whether the CC induced amelioration in body weight recovery was associated with reduced epithelial damage and immune cell infiltration, we assessed epithelial damage and infiltration in the large intestine by histology. Mice treated with PBS, CM, or SM showed a disruption of the epithelium and clear signs of infiltration (Figure 5d, Supplementary Table S6). In contrast, CC or FMT treated animals showed milder epithelial damage and infiltration, closer to the H2O group that did not receive DSS than to the PBS group. This supports our hypothesis that CC and FMT alleviated intestinal injury from DSS (Figure 5d,e; Supplementary Tables S7,S8). The mice treated with the CC and FMT also had reduced shortening of the colon and less pronounced increase in spleen weight when compared to the other DSS treated animals (Supplementary Figure S5, Supplementary Tables S9,S10). These results were confirmed in an independent repetition of the experiment (Supplementary Figure S6, Supplementary Tables S11-15) and suggest that only the CC matched the beneficial effect of FMT on the DSS-induced disease.

## Discussion

Functionally designed microbial consortia are being heralded as the next generation of microbiome-based therapeutics to be able to safely replace fecal transplants and feces-based products^13^. So far, only few approaches to rational consortium design based on function have been rep orted^44-46^. To our best knowledge, a solution to the biotechnological problem of producing consortia that capture the key properties of fecal transplants in a scalable, robust, and cost-efficient manner has remained elusive. Here, we present an approach for both bottom-up functional design and subsequent production of consortia using co-cultures. In this setting, producing consortium LBPs through co-culturing has important advantages compared to individually culturing and subsequently mixing strains, both on the functional and technological side.

Co-culturing enabled the *in vitro* growth of typically hard to culture strains at yields that are relevant for production. Determining the right conditions to cultivate many intestinal bacteria in the laboratory is a challenge in itself ^47^, and amount of resources required to further optimize the conditions to achieve acceptable yields are often prohibitive. Our results show that the function-based design approach produces consortia that can grow and stably coexist over extended periods of time in common cultivation conditions. This greatly streamlines any downstream optimization when moving from lab-scale to production-scale. Furthermore, the biomass yields of many individual strains is up to ten-fold higher in co-culture than monocultures despite the expectation that strains compete for a primary energy source. Hence, co-culturing a trophically interacting consortium both simplifies technological translation and improves biomass yields.

Co-culturing also better represents the *in situ* growth environment than individual monocultures. Both bacterial growth and specific metabolic activity depend on the environmental context, with respect abiotic parameters such as temperature, pH, redox, growth substrate, etc., as well as biotic parameters like competitive or positive interactions with coexisting strains^48^. Our results show that co-culturing preconditions the strains for rapid and balanced growth. In contrast, mixes of individually cultured strains have long lag phases while they adapt to new culture conditions. This is especially important when considering microbiome interventions in patients, where the processes of bacterial growth and washout due to transit contribute to the success or failure of bacterial engraftment. While no *in vitro* growth environment will fully capture all the intricacies of the human intestine, integrating part of the bacterial biotic environment into the production environment can better prime the consortium for growth after administration.

Finally, producing consortia using co-culturing requires only one single biotechnological production process that weakly depends on the number of strains. In contrast, when strains are produced in monoculture and subsequently mixed, increasing the number of strains to tens or even hundreds implies an increase in the required production equipment and infrastructure that is directly proportional in the number of strains. Co-culturing offers an elegant solution to this problem, independent of the number strains in the LBP.

This study puts forward a blueprint for the bottom-up design and subsequent production of defined consortia as LBPs to replace FMT or elicit specific microbiome modulations. Our approach to developing LBPs is based on a combination of ecological and biotechnological perspectives. On the ecological side, strain selection incorporates the metabolic dependencies required for a consortium-level functional output. Here, the explicit measurements of the metabolic activities *in vitro* are essential, as genome-based *in silico* methods could only predict a small subset of their actual metabolic capabilities. On the biotechnological side, continuous co-cultivation is the mere *in vitro* equivalent of growth in the intestine, incorporating flow, and abiotic and biotic environments. This resulted in a synthetic LBP that was able to match the therapeutic effect of FMT in an acute DSS model of colitis in mice, while a simple mix of the same strains was not. This implies that the production process imparts properties onto the consortium in a product-by-process manner. We expect that the co-cultivation of consortia will provide an efficient and highly controlled solution for LBP production adapted to GMP conditions. Based on our results, we believe that our approach will support the evolution to the next generation of microbiome products.

## Acknowledgements

PharmaBiome AG and ETH Zurich provided infrastructure and financial support. We thank Christophe Chassard for support with the strain selection, Markus Reichlin, Carmen Menzi, Florian Rosenthal, Marco Meola for their support in experiment planning and analytics. PharmaBiome AG and ETH Zurich were supported by Innosuisse Innovation Projects 40713.1 IP-LS and 27873_1 PFLS-LS.

## Author Contributions

F.K., L.A., M.R.S., G.R., C.L., and T.W conceptualized the project. F.K., G.E.L., M.R.S., G.R., C.L., T.W planned the experiments. F.K, L.A., and T.W. developed methods. M.R.S. planned and performed mouse experiments. F.K. and L.A. assisted with mouse experiments. F.K. and M.R.S. performed analyses. G.E.L. and PB. analyzed and curated data. T.W, C.L., and G.E.L. supervised the work. F.K., G.E.L., C.L., and T.W. wrote the original draft. All authors reviewed and edited the manuscript.

## Methods

### Assignment of essential reactions and strain selection

We assigned the essential metabolic reactions to a panel of intestinal bacterial strains. We measured their metabolic activity by cultivation on five different media: 1) Yeast extract, casitone, fatty acids medium (YCFA); 2) Glucose, soluble starch, cellobiose medium (M2GSC); 3-5) variants of M2GSC, where the ‘GSC’ carbon sources were exchanged by the intermediate metabolites lactate (M2LT), formate (M2FO), and succinate (M2SU), respectively Cultivation was strictly anaerobic in Hungate tubes. We assigned reactions A1-A6 to the strains based on media 1 and 2, and the reactions B1-B5 based on growth in media 3-5 (Supplementary Table S3). Growth was assessed after 48 h on media 1 and 2 and after 48 h and 7 days on media 3-5 to account for typically slow utilization rates for intermediate metabolites.

The YCFA and M2GSC media were prepared as described in the refs^49,50^. M2SU, M2LT, and M2FO contained 80 mM of sodium succinate (SU), 50 mM of DL-lactate (LT), or 60 mM of formate (FO), respectively. All media components were supplied by Sigma-Aldrich (Buchs, Switzerland) unless otherwise noted. We further supplemented the media with a redox potential indicator (resazurin, 1 mg/L). Liquid media were boiled, flushed with 100% O2-free CO2, dispensed into CO2-flushed Hungate tubes, sealed with butyl rubber septa (Bellco Glass, Vineland, USA), and autoclaved before use.

For the cultivation of pre-cultures, 0.5 mL of cryopreserved isolates (1 mL of 48 h old culture mixed with 1 mL of respective fresh medium containing 60% of glycerol, stored at −80 °C) were inoculated in Hungate tubes containing 8 mL of the respective preferred growth medium of the strain (see Supplementary Table S3). Optical density measurements confirmed growth. Subsequently, 0.1 mL of 48 h old single cultures were used to inoculate 8 mL of each of the five cultivation media in duplicates. To assign the respective A or B reactions, consumption and/or production of fatty acids was measured using high-pressure liquid chromatography with refractive index (HPLC-RI).

Literature data on hydrogen utilization or oxygen tolerability were used to assigned C-reactions to the strains, whereby oxygen reduction was assigned to strains reported to grow under aerobic or microaerophilic conditions and hydrogen consumption to acetogenic or methanogenic gut bacteria^41-43^.

### Preparation of single strains for inoculation

For the inoculation of bioreactors, strain pre-cultures were grown as described above. Equal volume inocula were produced by mixing 0.5 mL of each of the pre-cultures. ‘Controlled ratios’ were chosen to mimic the natural abundance of the respective genera in the host-microbiota from which seven out of the nine strains were isolated. To additionally improve the competitiveness of slow-growers and prevent growth inhibition through inhibitory concentrations of intermediate metabolites, the ratio of fast-growing primary degrader *Lb. rhamnosus* was diluted tenfold, and the ratios of secondary degraders *P.faecium, E. limosum*, and *A. lactatifermentans* were increased ten-fold compared to their relative abundance in the donor profile. The final concentrations are listed in Supplementary Table S5.

### Co-culture fermentation medium, PBMF009

We designed a growth medium—PBMF009—based on YCFA and M2GSC and adapted it to contain a maximal amount of complex carbohydrates and minimal amount of animal-derived ingredients (Supplementary Table S1). We did not include meat extract or rumen fluid. The carbon sources consisted of cellobiose (1.5 g/L), Fibru-lose F97 (1 g/L, Cosucra-Group Warcoing SA, Warcoing, Belgium), soluble potato starch (1.5 g/L) and resistant pea starch N-735 (2 g/L, Roquette Freres, Lestrem, France). For Hungate tube and serum flask experiments, we additionally buffered 3x to prevent rapid acidification of the culture (3xb PBMF009, Supplementary Table S1).

### Hungate tube cultivations

For individual strain cultures and co-cultures, we took 0.1 mL of 48 h-old single cultures or 0.1 mL of a mix of ‘equal volumes’ of all nine strains, respectively, and inoculated Hungate tubes containing 8 mL of 3xb PBMF009 medium. The tubes were then incubated for 48 h at 37 °C. We confirmed growth with optical density, measured metabolites with HPLC-RI, and quantified growth with qPCR at times t = 0 and t = 48 h. We performed three independent biological replicates of the cultivations with three technical replicates per condition.

### Bioreactor cultivations

Controlled fermentations were carried out in 0.3 L (Sixfors) or 1 L (Multifors) flat bottom glass bioreactors (Infors AG, Bottmingen, Switzerland), or 0.5 L Sartorius qPlus microbial version bioreactors (Sartorius AG, Gottingen, Germany). To maintain anaerobiosis, bioreactors were continuously flushed with CO2. The redox potential was monitored using EASYFERMPLUS ORP ARC probes (Hamilton, Bonaduz, Switzerland) and the supplier’s software HDM (Version 1.0; Hamilton). Medium feed, temperature, pH, stirring speed, and base consumption were monitored throughout the fermentation using the supplier’s software IRIS V5 (Version: 5; Infers AG), or DCU (Sartorius AG). pH was maintained by the automatic addition of 2.5 M NaOH.

Batch fermentation bioreactors were inoculated at 0.33% v/v and fed-batch fermentation bioreactors at 0.67% v/v (of the initial volume) with the respective mixing ratio of the nine consortium strains to balance for the medium refill during the fed-batch fermentation. Fermentation conditions were chosen to reflect conditions of the human colon and prevent a possible washout of slow growing strains, at 37 °C, pH = 6.0, and a mean retention time of 50 h (D = 0.02 h^−1^). Tie bioreactors were initialized with either batch or fed-batch mode for the first 48 h. In fed-batch mode, the first half of the fermentation medium was added at t = 0 and the second half was added at t = 24 h of fermentation. After 48 h, bioreactors were set to continuous mode. Bioreactor effluent samples were taken daily unless otherwise noted, and strain composition was measured using qPCR and metabolites by HPLC-RI.

### Reactivation in serum flasks

To assess whether the metabolic and compositional balance of the continuously co-cultured nine strain mix persists after a reactivation step, we performed a batch fermentation experiment in serum flasks and compared the regrowth of the CC with a mixture of individually grown single strains after inoculation. To produce the two different inocula, strains were either cultured individually in Hungate tubes for 48 h in their preferred growth medium (Supplementary Table S3) or as a co-culture in a controlled-ratio/fed-batch/continuous bioreactor as described in the previous section. On day ten of co-cultivation, when equilibrium was reached, we determined the abundance of each strain by qPCR. At the same time, viable cells in the Hungate tube cultures were measured by flow cytometry using live/dead staining approach (see protocol below). All measurements were performed in triplicates. For inoculation of the serum flasks, the individually cultured strains were mixed to match the absolute abundance of strains in the co-culture from the bioreactor. Batch fermentations were inoculated in triplicates with 0.1 mL of either of the two inocula in 10 mL of 3xb PBMF009 medium and were incubated and shaken (120 rpm, Celltron, Infers AG) at 37 °C for 48 h. We measured growth by optical density, pH, consortium composition by qPCR, and metabolites by HPLC-RI every 12 h.

### Quantitative PCR analysis

DNA extraction of samples was performed according to an adapted version of the “DNA purification from human feces using the Maxwell® RSC instrument and the Maxwell® RSC PureFood GMO and Authentication Kit” developed by Promega (Promega Corporation, Dübendorf, Switzerland). Instead of using 250 mg of fecal matter for the extraction process, we directly added 1 mL of CTAB Buffer to the pellet of a 1 mL liquid culture. After heating the samples for 5 min at 95 °C we transferred the liquid to a 2 mL Lysis Matrix E tube (MP Biomedicals, Lucerne, Switzerland) and bead beated 2-times for 40 sec at 6 m/sec using a FastPrep™ 24 5G Bead Beating System (MP Biomedicals). After the bead beating step, we centrifuged the samples at RT for 10 min at 14’000 g and transferred the supernatant to a new 2 mL Eppendorf tube and proceeded with the supplier’s guidelines. Quantitative real-time PCR (qPCR) was performed by using an ABI 7000 Sequence Detection System apparatus with 7000 system software version 1.2.3 (Applied Biosystems) or a magnet induction cycler (MIC) from Biomolecular systems (Labgene Scientific SA, Chatel-St-Denis, Switzerland). qPCR to enumerate 16S rRNA gene copies was performed by mixing one pL of sample genomic DNA with 2x Kapa Sybr Fast qPCR Mastermix (Biolabo Scientifics Instruments, Basel, Switzerland) in a total volume of 25 pL. We used genus-specific primers (Microsynth AG; Balgach, Switzerland) and validated no cross-reactivity on genomic DNA extracts of all other consortium members (Supplementary Table S2). Amplifications were performed with the following temperature profile: 1 cycle at 95 °C for 10 min to denature DNA and activate the polymerase, followed by 40 cycles of 95 °C for 30 sec, 60 °C for 1 min. A dissociation step was added to control the amplification specificity. A melting curve analysis was carried out to ensure the specificity of the amplification products. For quantification, 10-log-fold standard curves ranging from 102 to 107 copies were produced using the full-length amplicons of the purified 16S rRNA gene (pGEM®-T Vector Systems, Promega) of each of the nine PB002 strains to convert the threshold cycle (Ct) values into the average cells of target bacterial genomes present in 1 mL of fermentation effluent. As a negative control water was used. Sample DNA was analyzed in triplicates and results were normalized by the number of 16S rRNA copies per strain (for exact numbers see Supplementary Table S2).

### Whole genome sequencing of strains

High molecular weight DNA was extracted using the Maxwell® RSC PureFood GMO and Authentication Kit as for qPCR (Promega). The high molecular weight DNA was sent to the Functional Genomic Center Zurich (FGCZ, Zurich, Switzerland) for PacBio Sequel SMRT Cell sequencing. Additionally, high quality DNA was sent to FGCZ for library prep using the 2 × 150 bp True Seq kit and Illumina MiSeq Sequencing. For the assembly, PacBio long-reads were assembled using Flye 2.4.4^51^. This assembly was used in the hybrid assembly with the Illumina short reads using Unicycler 0.4.4^52^. The resulting assembly was annotated using Prokka 1.13.3^53^. We predicted the metabolic activities of the whole genomes with gutSMASH^54^.

### Metabolic analysis using HPLC-RI

The concentrations of SCFAs (formate, propionate, acetate, butyrate, and valerate), branched-chain fatty acids (BCFAs) (isobutyrate and isovalerate), and intermediate metabolites (lactate and succinate) were measured using HPLC-RI. Cultures of 1 mL or 100 mg of mouse cecal content homogenized in 600 pL of HPLC eluent were centrifuged (14,000 g, 10 min, 4 °C) and subsequently filtered into 1.5 mL short thread vials with crimp caps (VWR International GmbH, Schlieren, Switzerland) using 0.2 μm regenerated cellulose membrane filters (Phenomenex Inc., Aschaffenburg, Germany). Analyses were performed with a VWR Hitachi Chromaster 5450 RI-Detector using a Rezex ROA-Organic Acid (4%) precolumn connected to a Rezex ROA-Organic Acid (8%) column, equipped with a Security Guard Carbo-H cartridge (4 × 3.0 mm). Volumes of 40 pL were injected into the HPLC with a flow rate of 0.6 mL/min at a constant column temperature of 80 °C using a mixture of H2SO4 (10 mM) and Na-azide (0.05 g/L), as eluent. Concentrations were determined using external standards by comparing the retention time (all compounds were purchased from Sigma-Aldrich). Peaks were integrated using the EZChromElite software (Version V3.3.2.SP2, Hitachi High Tech Science Corporation). Tie limit of detection was defined as >0.8 mM for succinate, lactate, and acetate or >0.5 mM for all other SCFAs.

### Viability measurement using live/dead staining

We measured the viability of the individually cultivated strains after 48 h of incubation with flow cytometry. **A** double staining assay with the two nucleic acid dyes SYBR Green (SG) and propidium iodide (PI) was used to differentiate between cells with intact and damaged cytoplasmic membranes, as described by Nevel *et al*. [55]. Samples were diluted 100-fold with filtered (0.20 μm) PBS to get an appropriate bacterial concentration for flow cytometry. Samples of 30 µL were stained either with 3 µL of SG (10 µL of SG stock solution from Life Technologies Europe BV, Zug, Switzerland, in 990 µL dimethylsulfoxide) or 3 µL of SG combined with PI (SGPI) (10 µL of SYBR Green stock solution and 20 µL of PI stock solution 20 mM from Life Technologies Europe BV, in 970 µL dimethylsulfoxide) and mixed by vortexing. Samples were stained for 20 min at RT protected from light. To determine bacterial cell numbers, 30 µL of Flow-CountTM Fluorospheres beads (Beckman Coulter, Nyon, Switzerland) at known concentrations were added to 30 µLof the diluted samples. Before measuring, the samples were again mixed by vortexing. Each sample was diluted, stained, and measured in triplicates. Each replicate was stained twice, once with SG only for total cell count, and once with SGPI for differentiating viable and dead cells. SGPI stained samples were always measured first to limit aerobic conditions and avoid false Pl-positive results because of cells dying during the incubation period.

### Dextran Sulfate Sodium-induced acute colitis mouse model

To assess the effect of PB002 administration on recovery from intestinal inflammation, the acute DSS-induced colitis mouse model was used. Female C57BL/6J mice were purchased from Janvier labs (Le Genest-Saint-Isle, France). Upon arrival at the animal facility of the University Hospital Zurich, the mice were acclimatized for two weeks before the start of the experiment. All mice were housed in a barrier-protected specific pathogen-free unit in individually ventilated cages, with a 12:12 hour light/dark cycle and an artificial light of approximately 40 Lux in the cage. The animals were kept under controlled humidity (45-55%) and temperature (21 ± 1°C). Mice had access to sterilized drinking water and to pelleted and extruded regular mouse chow (diet 3436., Kliba Nafag, Switzerland) *ad libitum*. Mice with a body weight between 20 g and 23 g were used for all experiments. 3% DSS (MP Biomedicals, Carlsbad, CA) was added to the drinking water for seven days (day 1 until day 8 of the experiment) to induce acute colitis. On day 8, the DSS-containing water was replaced with normal water, and mice were randomly allocated to control and experimental groups to obtain groups with equal body weight at the treatment start. Each treatment group consisted of 4-5 mice. The control (sterile PBS) and treatment substances were administered daily per oral gavage (200 pL) on days 8, 9, and 10. The treatment substances were prepared as follows:

The strain mix was prepared akin to the ‘controlled ratio’ inoculum. ‘Conditioned medium consisted of the centrifuged (10,000 g, 10 min, 4 °C) and subsequently filtered (0.45 μm Nylon filter) supernatant of the CC after eight days of continuous co-cultivation from a ‘controlled ratio’ inoculation taken from the same bioreactor, and at the same time as for PB002. To produce Human FMT, a fecal suspension was prepared by adding 100 g fecal sample of a healthy male donor (approved by the ethical commission of Zurich, Switzerland; Kantonale Ethikkomission Zü rich, BASEC-Nr. 2017-01290) to 500 mL sterile saline. The suspension was homogenized in a blender for 2 min. Particles were then allowed to sediment for 20 sec. Particulate material was removed by passing the slurry through a sterile gauze. The slurry was centrifuged (6,000 g, 15 min, 4 °C), and the supernatant was subsequently removed. The pellet was resuspended in 15.3 mL sterile saline and filtered with a Falcon Cell Strainer 100 μm Nylon Filter (VWR, Dietikon, Switzerland) to remove particles. Glycerol was added to reach a final concentration of 10% (as specified by Youngster *et al*. [56]). The FMT samples were frozen at −80 °C and thawed on ice 1 h before use.

Body weight was monitored daily and mice were euthanized on day 16 to collect colon and spleen specimens, which served as a proxy for colitis severity. Moreover, bacterial metabolites of cecal content were analyzed using HPLC-RI. Animal experiments were carried out according to Swiss animal welfare laws and were approved by the veterinary authorities of Zurich, Switzerland (Veterinaramt des Kantons Zurich, approval no. ZH255/2014 and ZH171/2017).

Hematoxylin and eosin staining and histological assessment of colitis severity Formalin-fixed, paraffin-embedded samples of the most distal 1.5 cm of the colon and the distal part of the cecum were cut into 5 μm sections, and stained with hematoxylin and eosin (H&E) according to standard procedures. In brief, formalin-fixed, paraffin-embedded tissue samples were deparaffinized in HistoClear for 2x 10 min, followed by incubation in a descending ethanol series (100%, 96%, 70% ethanol). Samples were briefly incubated in double-distilled H2O prior to incubation in hematoxylin for 10 minutes and subsequently rinsed with tap water before differentiation in a solution consisting of 20% HC1 and 80% ethanol for twice one second. Samples were then rinsed for 10 min with tap water and incubated for 15 seconds in a 1% eosin solution. Samples were dehydrated in an ascending ethanol series, incubated in HistoClear, and finally mounted. The slides were analyzed using an AxioCam HRc (Zeiss, Jena, Germany) on a Zeiss Axio Imager.Z2 microscope (Zeiss) and images captured using the Axio-Vision Release 4.8.2 software (Zeiss). For colitis severity assessment, the following score was applied: score for epithelial damage: 0, normal morphology; 1, loss of goblet cells; 2, loss of goblet cells in large areas; 3, loss of crypts; 4, loss of crypts in large areas. The score for infiltration: 0, no infiltrate; 1, infiltrate around crypt base; 2, infiltrate reaching to *L. muscularis mucosae-*, 3, extensive infiltration reaching the *L. muscularis* and thickening of the mucosa with abundant edema; 4, infiltration of the *L. submucosae*. The total histological score represents the sum of the scores for epithelial damage and infiltration. Scoring activities were performed in a blinded manner.

### Statistical Analysis

Graphing and statistical analysis were performed in R version 4.0.2^57^. qPCR data were normalized by dividing by the number of 16 S rRNA gene copies per strain. Principle component analysis (PCA) was performed on the relative abundances of the consortium strains after a center log-ratio transformation. The correlation between metabolites was computed as the Pearson correlation between the individual metabolite time series. Mouse weights were normalized by their weight at day 0. The area under the curve (AUC) was computed as the sum of relative mouse weights throughout the experiment. Comparisons of readouts between mouse treatment groups were done using a linear model with the PBS group as the intercept.

## Supplementary Figures

**Figure S1:**
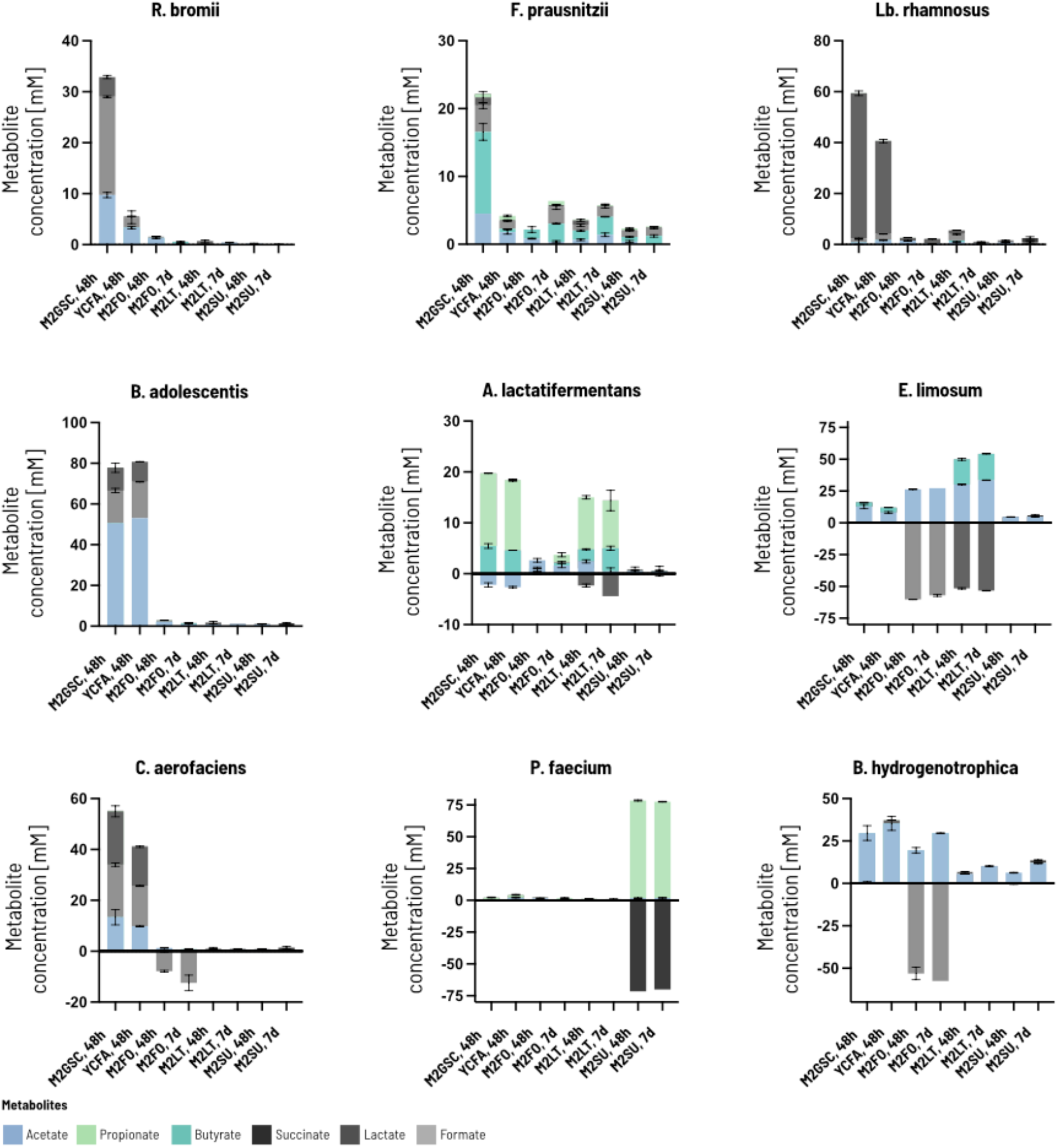
Metabolic profile of the selected nine strains as single cultures in five different cultivation media in Hungate tubes. The produced or consumed metabolites were used to assign A, B and C reactions to the respective single strains. Metabolite concentrations were quantified by HPLC-RI after 48 h and / or 7 days of batch incubations. Strains were inoculated in a 1:100 ratio. Data shown for two technical replicates per medium and strain as means + SD.

**Figure S2:**
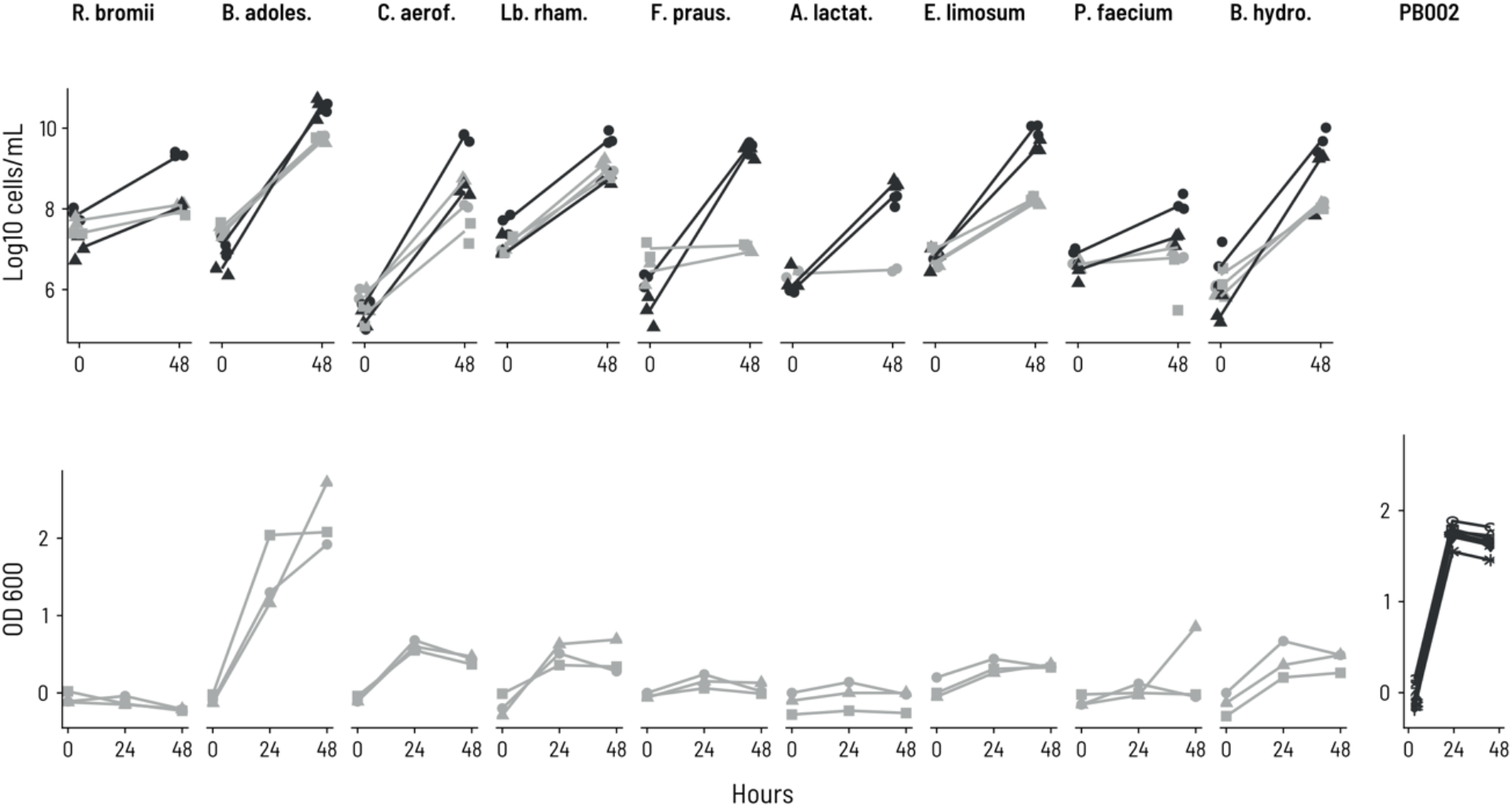
Growth of strains in single culture vs. co-culture. Absolute growth of the selected nine PB002 strains inoculated either as single strains or as co-cultivated consortium as quantified by qPCR after 48 h of batch incubations in Hungate tubes on 3xb PBMF009 medium. The top panel shows cell counts estimated by qPCR for 2 - 3 repetition experiments. Some measurements were left out due to failure of DNA extraction. Four of the nine strains showed no or only weak growth in monoculture on the 3xb PBMF009 medium. In contrast, co-culturing enabled the growth of all nine strains of PB002. The bottom panel shows the OD_600_ measurement of the same single strain cultures. OD 600 measurements of the co-culture is on the right. Light grey: single stains, dark grey: co-culture. Circles, squares, and triangles indicate three replicates. qPCR data is normalized by the 16S rRNA gene copy number of each strain.

**Figure S3:**
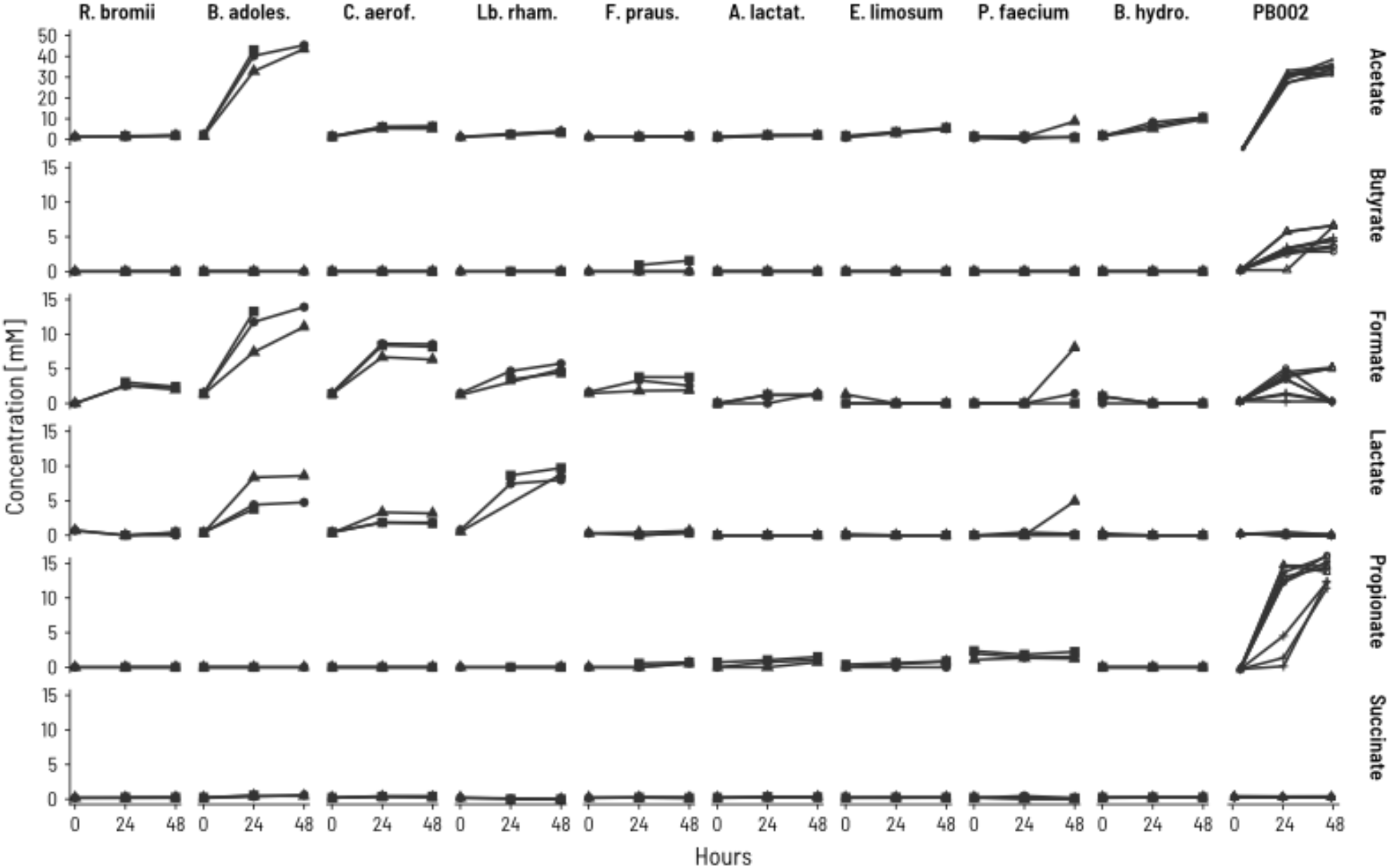
Metabolites produced in single cultures and co-cultures in Hungate tubes. Metabolite concentrations of the selected nine PB002 strains inoculated either as single strains or as co-cultivated consortium (PB002)as quantified by HPLC-RI after 0 h, 24 h, and 48 h of batch incubations in Hungate tubes on PBMF009 medium. Circles, squares, and triangles indicate three biological replicates.

**Figure S4:**
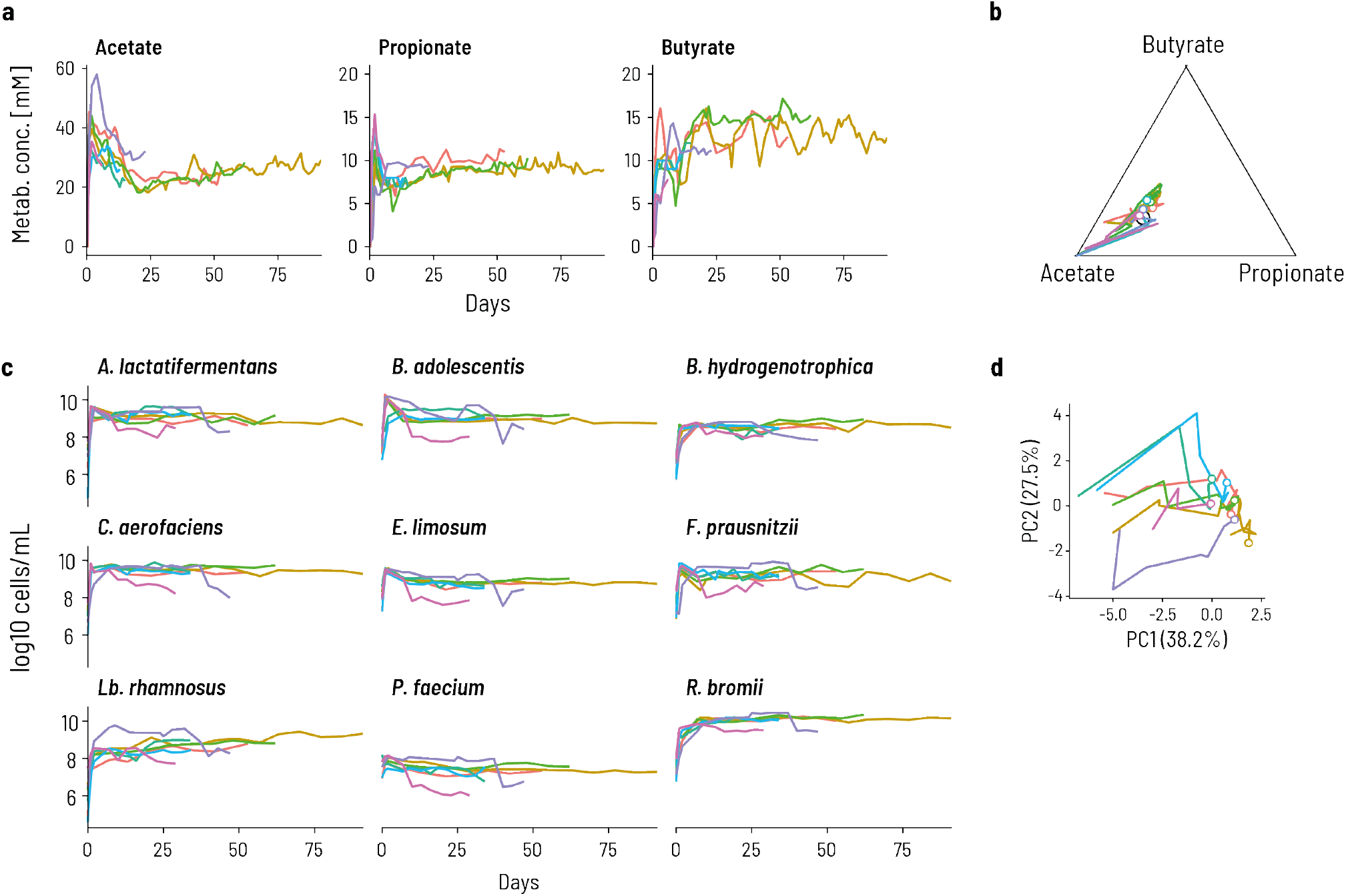
A repeatable and robust metabolic profile and community composition is reached within 11 days of continuous cocultivation that follows an initial 48 h of fed-batch fermentation, **a**. The metabolite profile contained exclusively end metabolites near the 3:1:1 ratio, reached the same concentrations in all repetitions, and remained stable for up to 92 days. **b**. The community composition also reached a common equilibrium and overall yield, despite early transient differences, as shown by absolute strain abundances measured by qPCR as well as the PCA plot. Data is shown for six replicate bioreactors, each in a different color. qPCR data are normalized by the 16S rRNA gene copy number of each strain.

**Figure S5:**
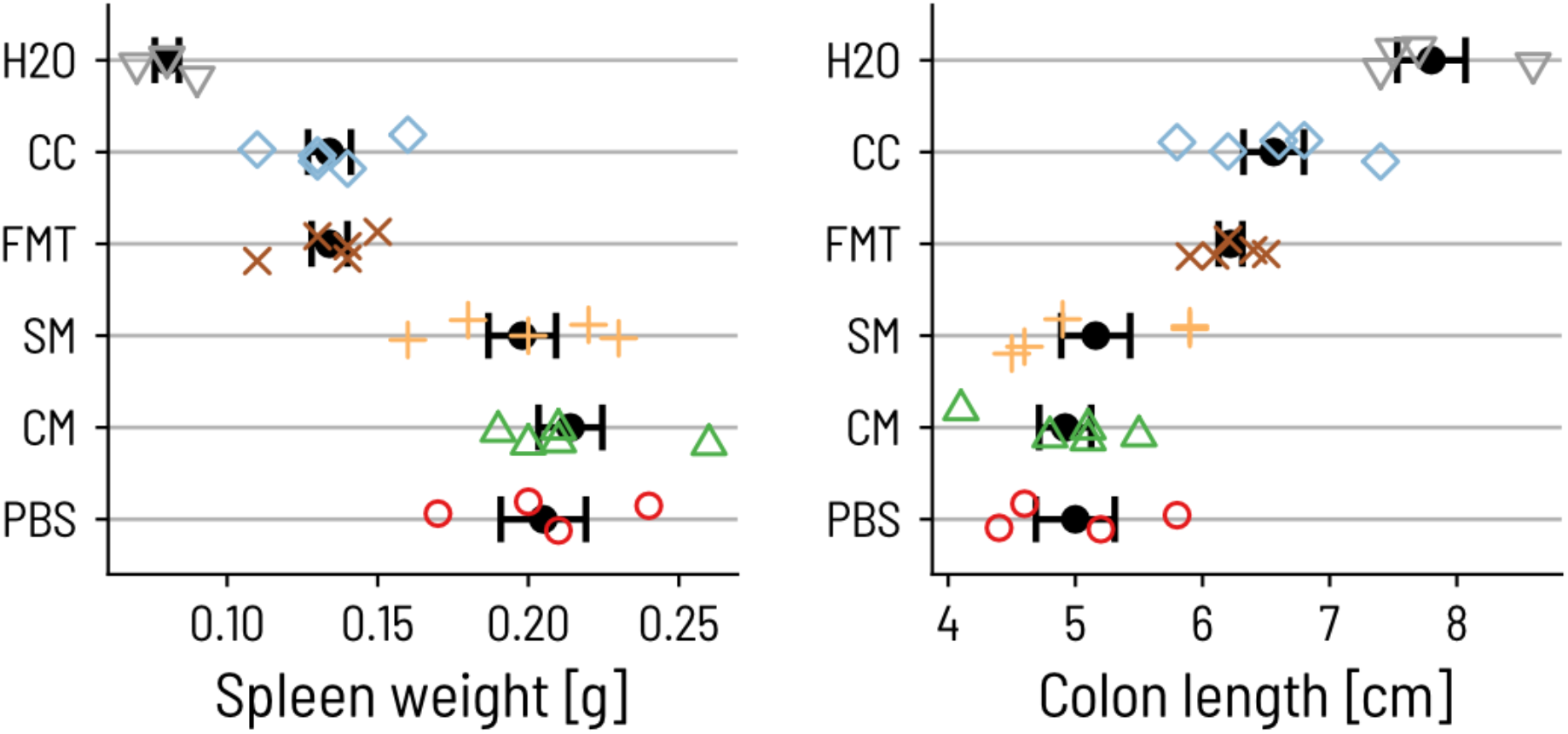
Spleen weight and colon length from the DSS mouse experiment. Mouse spleen weight (left panel) and colon length (right panel) of individual C57/B6 mice at day 16 of experimentation. Each symbol corresponds to one mouse. Error bars show the computed confidence interval for the mean.

**Figure S6:**
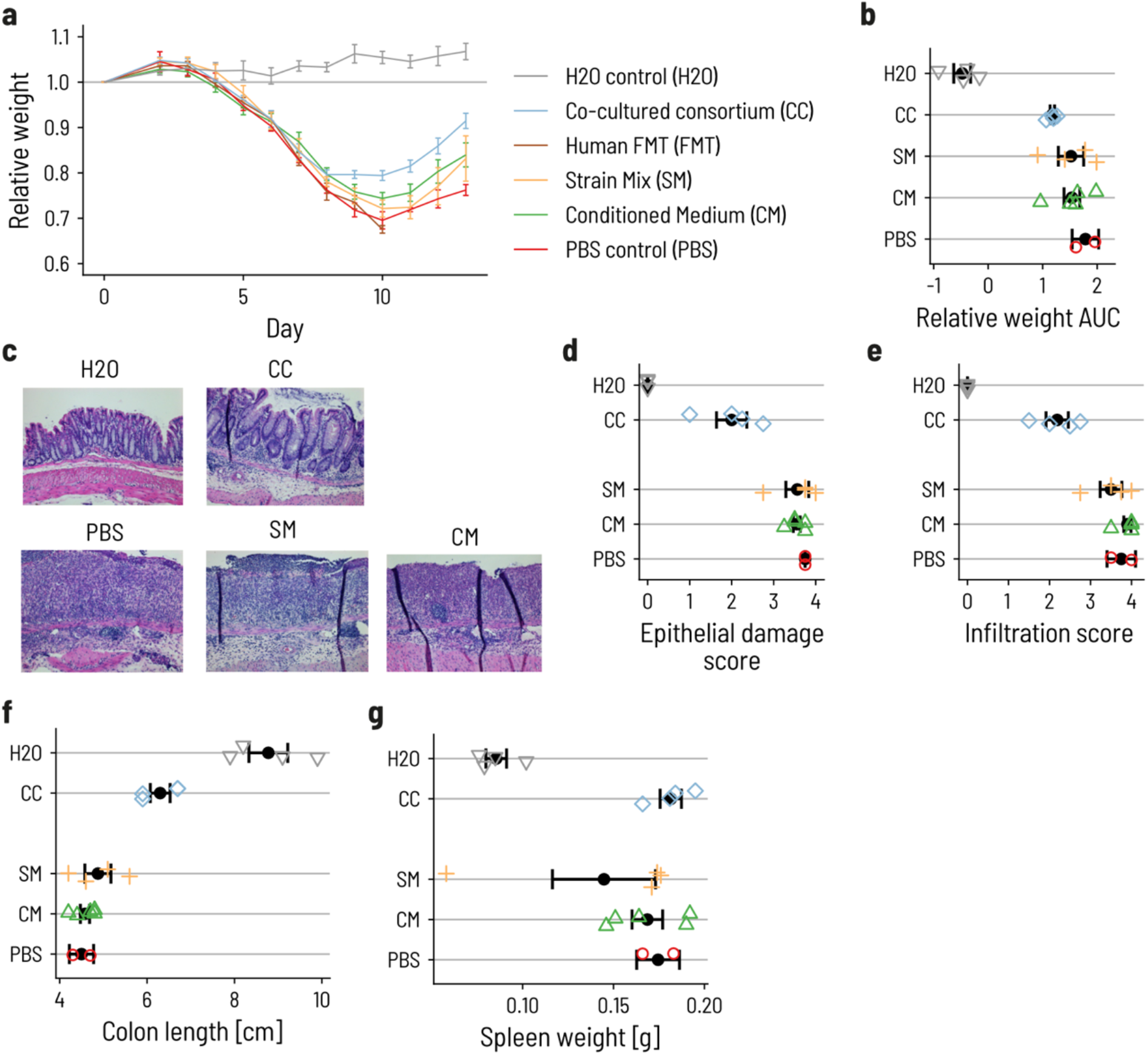
Repetition of the DSS mouse experiment, **a**. Relative weight loss/gain on average per group with error bars signifying the standard error (SE). Treatment group, receiving the CC, showed a weight recovery superior to all other treatment groups, **b.** Area under the curve (AUC) of the experiment, ***c***. Representative histological preparations of the large intestine on the day of sacrifice using H&E staining procedure. CC is the only treatment group showing structural recovery of the epithelium comparable to the healthy control group on the day of sacrifice. CM, SM, and PBS control still display substantial degradation and inflammation of the cecum’s epithelium, **d-e**. Histological assessment of the epithelial damage ***(d)*** and infiltration **(e)** of the distal colon at day 16. **f-g**. Colon length **(f)** and spleen weight ***(g)*** at day 16. All mice from the FMT treatment group were euthanized before the end of the experiment, due to weight loss > 30%. Data is shown for four mice per control groups and five mice per treatment groups.

## Supplementary Tables

**Table S 1:** Composition of nutritive growth medium PBMF009 (g/L) on the left side and the three-times buffered version of PBMF009 used in Hungate tube experiments on the right side.

| Constituent and supplier | PBMF009 | 3xb PBMF009 |
| --- | --- | --- |
|  | g/L | g/L |
| Amicase, casein acid hydrolysate, from bovine milk | 5 | 5 |
| Yeast extract for biotechnology Fermtech® | 2.5 | 2.5 |
| Cellobiose | 3 | 3 |
| Fibrulose F97 | 2 | 2 |
| Soluble potato starch | 3 | 3 |
| Resistant starch "Amidon de pois N-735" | 4 | 4 |
| Hemin | 0.025 | 0.025 |
| Resazurin | 0.001 | 0.001 |
| D-(+)-Biotin | 0.00001 | 0.00001 |
| Cobalamin | 0.00001 | 0.00001 |
| P-aminobenzoic acid | 0.00003 | 0.00003 |
| Folic acid | 0.00005 | 0.00005 |
| Pyridoxamine | 0.00015 | 0.00015 |
| K<sub>2</sub>HPO<sub>4</sub> | 0.15 | 0.066 |
| KH<sub>2</sub>PO<sub>4</sub> | 0.15 | 0.066 |
| NaCl | 0.3 | 0.132 |
| (NH<sub>4</sub>)SO<sub>4</sub> | 0.3 | 0.132 |
| MgSO<sub>4</sub> | 0.03 | 0.0132 |
| CaCl<sub>2</sub> | 0.03 | 0.0132 |
| Cysteine HCl | 1 | 1 |
| NaHCO<sub>3</sub> | 4 | 8 |

All Sigma-Aldrich (Buchs, Switzerland), except Fibrulose F97 from Cosucra-Group Warcoing SA (Warcoing, Belgium), and resistant starch "Amidon de pois N-735" from Roquette Frères (Lestrem, France) Abbreviation: 3xb = three-times buffered

**Table S 2:**
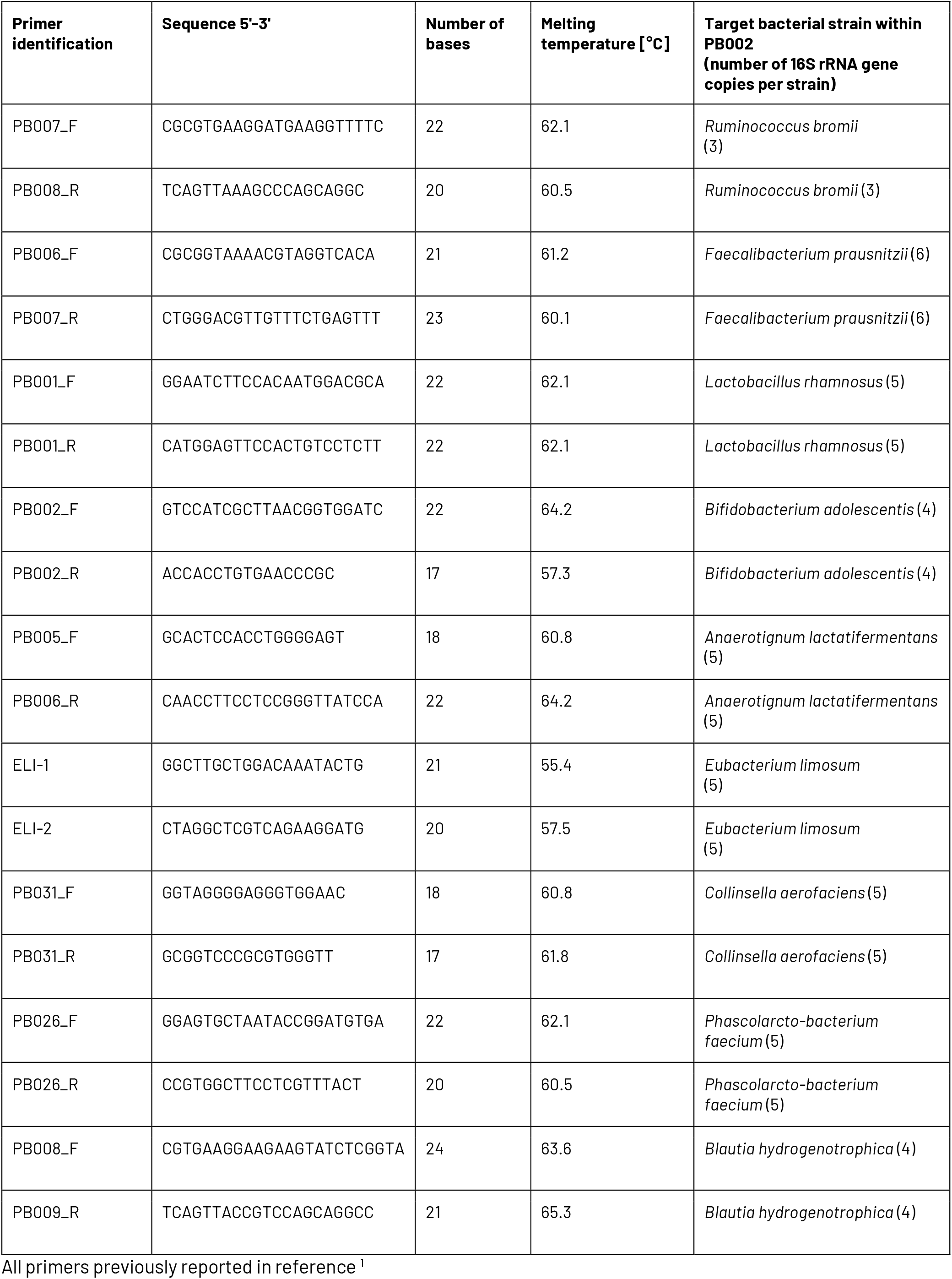
Genus specific primers used for the enumeration of the PB002 consortium members in fermentation effluent samples by qPCR analysis.

**Table S 3:**
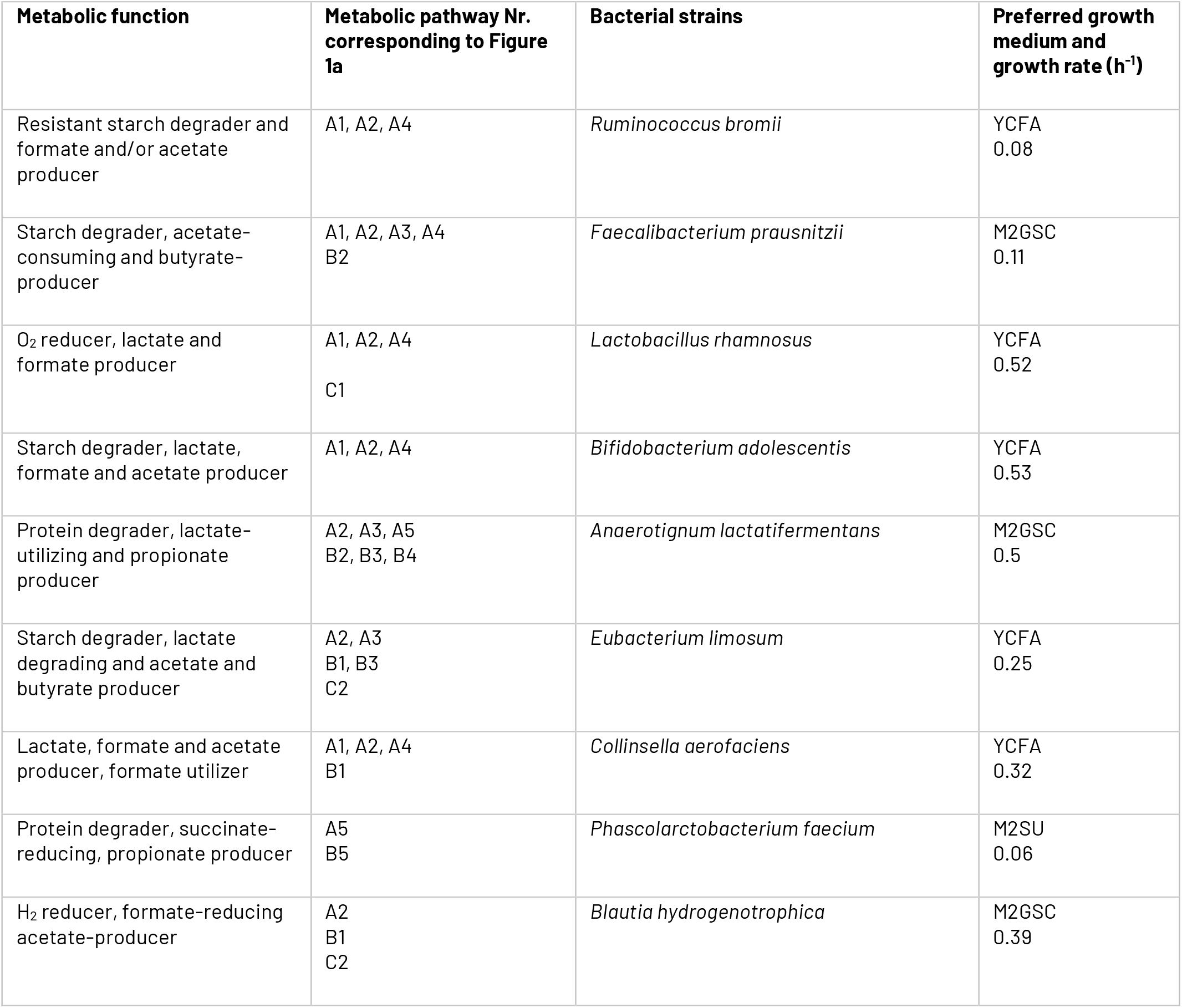
Metabolic functions and pathways affiliation of PB002. The metabolic pathway numbers according to Fig. 1a were assigned to the nine strains using Hungate tube batch fermentations on five different cultivation media. The preferred growth medium was defined as the medium in which the respective strains reached the highest OD_600_ within 48 h of batch fermentation. The specific growth rates were performed in the preferred growth medium.

**Table S 4:**
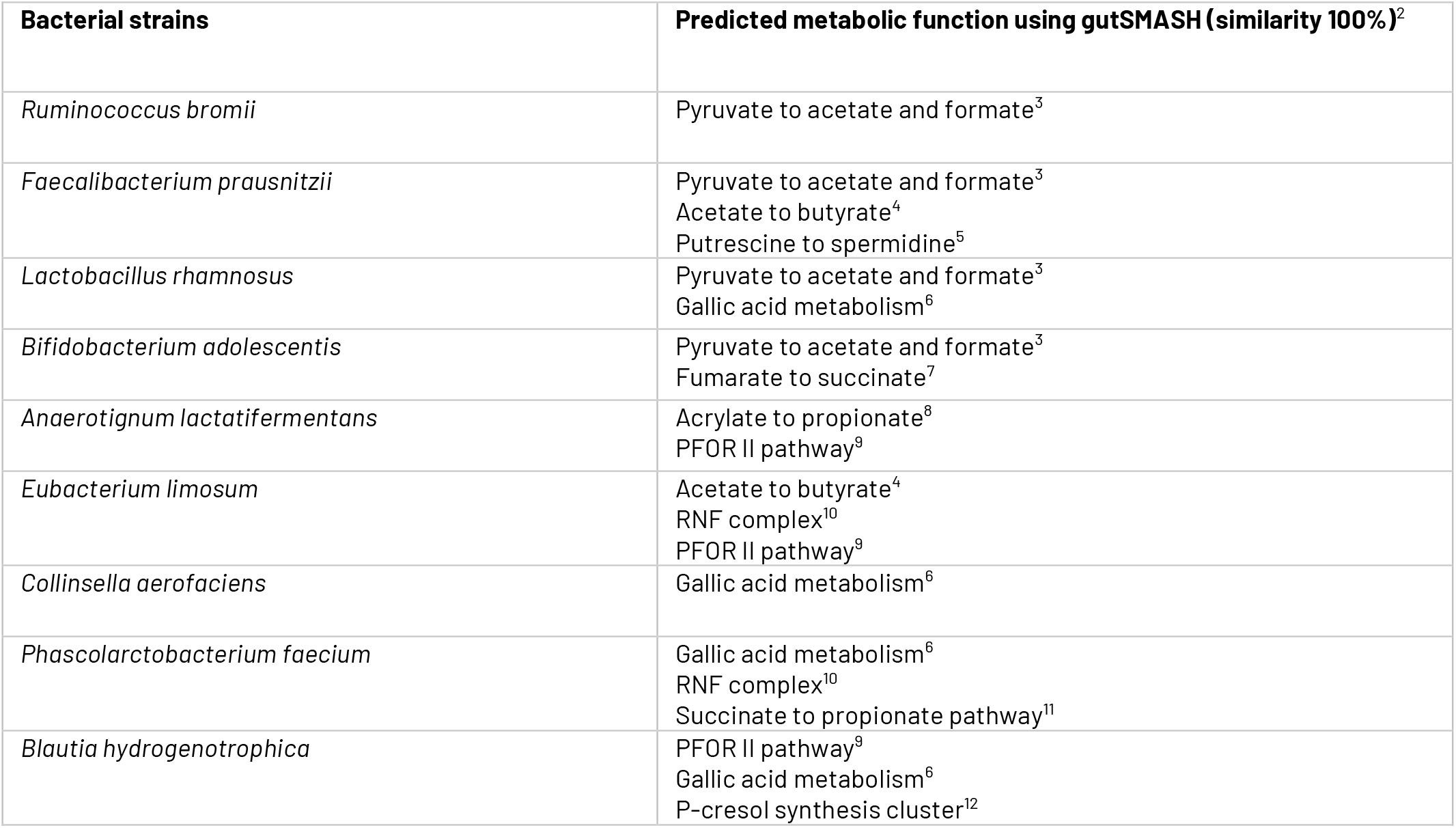
*In sili*cometabolic pathway affiliation. Metabolic pathways were predicted using gutSMASH with the whole genomes of the PB002 strains.

| Bacterial strains | Predicted metabolic function using gutSMASH (similarity 100%)<sup>2</sup> |
| --- | --- |
| *Ruminococcus bromii* | Pyruvate to acetate and formate<sup>3</sup> |
| *Faecalibacterium prausnitzii* | Pyruvate to acetate and formate<sup>3</sup><br>Acetate to butyrate<sup>4</sup><br>Putrescine to spermidine<sup>5</sup> |
| *Lactobacillus rhamnosus* | Pyruvate to acetate and formate<sup>3</sup><br>Gallic acid metabolism<sup>6</sup> |
| *Bifidobacterium adolescentis* | Pyruvate to acetate and formate<sup>3</sup><br>Fumarate to succinate<sup>7</sup> |
| *Anaerotignum lactatifermentans* | Acrylate to propionate<sup>8</sup><br>PFOR II pathway<sup>9</sup> |
| *Eubacterium limosum* | Acetate to butyrate<sup>4</sup><br>RNF complex<sup>10</sup><br>PFOR II pathway<sup>9</sup> |
| *Collinsella aerofaciens* | Gallic acid metabolism<sup>6</sup> |
| *Phascolarctobacterium faecium* | Gallic acid metabolism<sup>6</sup><br>RNF complex<sup>10</sup><br>Succinate to propionate pathway<sup>11</sup> |
| *Blautia hydrogenotrophica* | PFOR II pathway<sup>9</sup><br>Gallic acid metabolism<sup>6</sup><br>P-cresol synthesis cluster<sup>12</sup> |

**Table S 5:** Mixing ratios of the single strains for the ‘controlled ratio’ inoculum were selected to mimic the natural abundance of the respective genera in the host-microbiota from which seven out of the nine strains were isolated, as defined by next-generation sequencing (MiSeq, Starseq).

| Bacterial strains | Relative Abundance in donor microbiota (genus level) [%] | Mixing ratio of strains for 'controlled-ratio' (CR) inoculum (%) |
| --- | --- | --- |
| *Ruminococcus bromii* | 8.15 | 4 |
| *Faecalibacterium prausnitzii* | 13.65 | 30 |
| *Lactobacillus rhamnosus* | 0.003 | 0.0004 |
| *Bifidobacterium adolescentis* | 3.30 | 2 |
| *Anaerotignum lactatifermentans* | 3.00 | 20 |
| *Eubacterium limosum* | 3.75 | 20 |
| *Collinsella aerofaciens* | 1.45 | 2 |
| *Phascolarctobacterium faecium* | 2.35 | 20 |
| *Blautia hydrogenotrophica* | 2.50 | 2 |

**Table S6:**
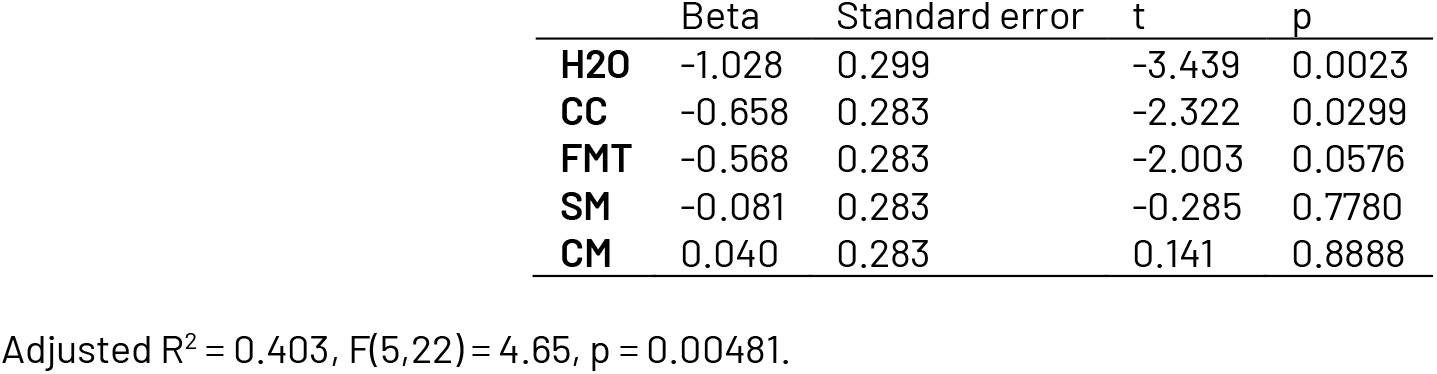
Regression coefficients for the weight area under the curve (AUC) on treatment compared to PBS for the first DSS experiment. A smaller AUC implies less weight loss.

**Table S7:** Regression coefficients for the epithelial damage score on treatment compared to PBS for the first DSS experiment. A larger score implies more damage.

|  | Beta | Standard error | t | p |
| --- | --- | --- | --- | --- |
| **H2O** | -3.625 | 0.334 | -10.849 | 2.69E-10 |
| **CC** | -1.825 | 0.317 | -5.757 | 8.61E-06 |
| **FMT** | -1.575 | 0.317 | -4.969 | 5.68E-05 |
| **SM** | -0.525 | 0.317 | -1.656 | 1.12E-01 |
| **CM** | -0.325 | 0.317 | -1.025 | 3.16E-01 |

Adjusted $R^2 = 0.859$ , $F(5,22) = 27.4$ , $p = 1.722 \times 10^{-6}$ .

**Table S8:** Regression coefficients for the infiltration score on treatment compared to PBS for the first DSS experiment. A larger score implies more damage.

|  | Beta | Standard error | t | p |
| --- | --- | --- | --- | --- |
| **H2O** | -3.625 | 0.2944 | -12.315 | 2.41E-11 |
| **CC** | -1.85 | 0.2793 | -6.625 | 1.16E-06 |
| **FMT** | -1.45 | 0.2793 | -5.192 | 3.31E-05 |
| **SM** | -0.55 | 0.2793 | -1.97 | 0.0616 |
| **CM** | -0.15 | 0.2793 | -0.537 | 0.5966 |

Adjusted $R^2 = 0.892$ , $F(5,22) = 45.4$ , $p = 7.33 \times 10^{-9}$ .

**Table S9:** Regression coefficients for the colon length on treatment compared to PBS for the first DSS experiment.

|  | Beta | Standard error | t | p |
| --- | --- | --- | --- | --- |
| **H2O** | 2.8 | 0.3931 | 7.122 | 3.84E-07 |
| **CC** | 1.56 | 0.373 | 4.183 | 0.000386 |
| **FMT** | 1.22 | 0.373 | 3.271 | 0.003492 |
| **SM** | 0.16 | 0.373 | 0.429 | 0.672085 |
| **CM** | -0.08 | 0.373 | -0.215 | 0.832128 |

Adjusted $R^2 = 0.760$ , $F(5,22) = 18.1$ , $p = 3.91 \times 10^{-7}$ .

**Table S10:** Regression coefficients for spleen weight on treatment compared to PBS for the first DSS experiment. A larger score implies more damage.

|  | Beta | Standard error | t | p |
| --- | --- | --- | --- | --- |
| **H2O** | -0.125 | 0.01591 | -7.856 | 7.97E-08 |
| **CC** | -0.071 | 0.0151 | -4.703 | 0.000108 |
| **FMT** | -0.071 | 0.0151 | -4.703 | 0.000108 |
| **SM** | -0.007 | 0.0151 | -0.464 | 0.6474 |
| **CM** | 0.009 | 0.0151 | 0.596 | 0.557115 |

Adjusted $R^2 = 0.813$ , $F(5,22) = 24.6$ , $p = 2.58 \times 10^{-8}$ .

**Table S11:** Regression coefficients for the weight area under the curve (AUC) on treatment compared to PBS for the second DSS experiment. A smaller AUC implies less weight loss.

|  | Beta | Standard error | t | p |
| --- | --- | --- | --- | --- |
| **H2O** | -2.2602 | 0.2927 | -7.723 | 2.06E-06 |
| **CC** | -0.6071 | 0.2927 | -2.075 | 0.0569 |
| **SM** | -0.2646 | 0.2927 | -0.904 | 0.3812 |
| **CM** | -0.2483 | 0.2827 | -0.878 | 0.3947 |

Adjusted $R^2 = 0.854$ , $F(4,14) = 27.4$ , $p = 1.72 \times 10^{-6}$ .

**Table S12:** Regression coefficients for the epithelial damage score on treatment compared to PBS for the second DSS experiment. A larger score implies more damage.

|  | Beta | Standard error | t | p |
| --- | --- | --- | --- | --- |
| **H2O** | -3.75 | 0.3819 | -9.82 | 1.17E-07 |
| **CC** | -1.75 | 0.3819 | -4.583 | 0.000426 |
| **SM** | -0.1875 | 0.3819 | -0.491 | 0.631019 |
| **CM** | -0.2 | 0.3689 | -0.542 | 0.596247 |

Adjusted $R^2 = 0.916$ , $F(4,14) = 50.4$ , $p = 3.70 \times 10^{-8}$ .

**Table S13:** Regression coefficients for the infiltration score on treatment compared to PBS for the second DSS experiment. A larger score implies more damage.

|  | Beta | Standard error | t | p |
| --- | --- | --- | --- | --- |
| **H2O** | -3.75 | 0.3372 | -11.123 | 2.46E-08 |
| **CC** | -1.5625 | 0.3372 | -4.634 | 0.000386 |
| **SM** | -0.25 | 0.3372 | -0.742 | 0.470645 |
| **CM** | 0.15 | 0.3257 | 0.461 | 0.652216 |

Adjusted $R^2 = 0.938$ , $F(4,14) = 69.4$ , $p = 4.52 \times 10^{-9}$ .

**Table S14:** Regression coefficients for the colon length on treatment compared to PBS for the second DSS experiment. A larger score implies more damage.

|  | Beta | Standard error | t | p |
| --- | --- | --- | --- | --- |
| **H2O** | 4.275 | 0.4955 | 8.628 | 5.62E-07 |
| **CC** | 1.8 | 0.4955 | 3.633 | 0.00272 |
| **SM** | 0.375 | 0.4955 | 0.757 | 0.46171 |
| **CM** | 0.08 | 0.4787 | 0.167 | 0.86967 |

Adjusted $R^2 = 0.893$ , $F(4,14) = 38.6$ , $p = 2.04 \times 10^{-7}$ .

**Table S15:** Regression coefficients for the spleen weight on treatment compared to PBS for the second DSS experiment. A larger score implies more damage.

|  | Beta | Standard error | t | p |
| --- | --- | --- | --- | --- |
| **H2O** | -0.089 | 0.02626 | -3.389 | 0.00441 |
| **CC** | 0.007 | 0.02626 | 0.267 | 0.79369 |
| **SM** | -0.02975 | 0.02626 | -1.133 | 0.27629 |
| **CM** | -0.0059 | 0.02537 | -0.233 | 0.81947 |

Adjusted $R^2 = 0.547$ , $F(4,14) = 6.43$ , $p = 0.00374$ .

